# A receptor-inactivation model for single-celled habituation in *Stentor coeruleus*

**DOI:** 10.1101/2024.11.05.622147

**Authors:** Deepa H Rajan, Wallace F. Marshall

## Abstract

The single-celled ciliate *Stentor* coeruleus demonstrates habituation to mechanical stimuli, showing that even single cells can manifest a basic form of learning. Although the ability of *Stentor* to habituate has been extensively documented, the mechanism of learning is currently not known. Here we take a bottom-up approach and investigate a simple biochemistry-based model based on prior electrophysiological measurements in *Stentor* along with general properties of receptor molecules. In this model, a mechanoreceptor senses the stimulus and leads to channel opening to change membrane potential, with a sufficient change in polarization triggering an action potential that drives contraction. Receptors that are activated can become internalized, after which they can either be degraded or recycled back to the cell surface. This activity-dependent internalization provides a potential means for the cell to learn. Stochastic simulations of this model confirm that it is capable of showing habituation similar to what is seen in actual *Stentor* cells, including the lack of dishabituation by strong stimuli and the apparently step-like response of individual cells during habituation. The model also can account for several habituation hallmarks that a previous two-state Markov model could not, namely, the dependence of habituation rate on stimulus magnitude, which had to be added onto the two state model but arises naturally in the receptor inactivation model; the rate of response recovery after cessation of stimulation; the ability of high frequency stimulus sequences to drive faster habituation that results in a lower response probability, and the potentiation of habituation by repeated rounds of training and recovery. The model makes the prediction that application of high force stimuli that do not normally habituate should drive habituation to weaker stimuli due to decrease in the receptor number, which serves as an internal hidden variable. We confirmed this prediction using two new sets of experiments involving alternation of weak and strong stimuli. Furthermore, the model predicts that training with high force stimuli delays response recovery to low force stimuli, which aligns with our new experimental data. The model also predicts subliminal accumulation, wherein continuation of training even after habituation has reached asymptotic levels should lead to delayed response recovery, which was also confirmed by new experiments. The model is unable to account for the phenomenon of rate sensitivity, in which habituation caused by higher frequency stimuli is more easily reversed leading to a frequency dependence of response recovery. Such rate sensitivity has not been reported in *Stentor*. Here we carried out a new set of experiments which are consistent with the model’s prediction of the lack of rate sensitivity. This work demonstrates how a simple model can suggest new ways to probe single-cell learning at an experimental level. Finally, we interpret the model in terms of a kernel estimator that the cell may use to guide its decisions about how to response to new stimuli as they arise based on information, or the lack thereof, from past stimuli.

## Introduction

Cells are much more than just amorphous bags of enzymes, and their structural complexity is mirrored by a startling level of behavioral complexity that one would normally associate with animals that have a nervous system (Jennings 1906; Vertosick 2002; Bray 2011; Baluska and Levin 2016; Larson 2023). In addition to carrying out decision-making processes, cells are able to carry out at least simple forms of learning (Tang and Marshall 2018). One of the most fundamental forms of learning, habituation, is defined as the reduction in response to a stimulus following repeated application of that stimulus (Harris 1943; Thompson and Spencer 1966; Groves 1970; Rankin 2009). Habituation has been extensively documented in the giant single-celled organism *Stentor coeruleus* (Jennings 1902; Wood 1970a; Wood 1970b; Wood 1988a; Wood 1988b; Rajan 2023a). *Stentor* is a giant ciliate most famous for its amazing powers of regeneration (Tartar 1961; Marshall 2021). When undisturbed, the cell stretches out into a cone-like shape, attaching to pond plants via a holdfast and feeding by means of a membranellar band of cilia at its anterior end (**Figure 1A**). When disturbed by a mechanical stimulus, the cell contracts downwards towards its point of attachment (Jones 1970; Newman 1972), by means of a series of parallel myonemes that run the length of the cell. These myonemes are composed of calcium binding EF hand proteins in the centrin family (Huang and Pitelka, 1973; Maloney 2005). The large size of *Stentor* cells, and the extensive shape change that occurs during contraction, makes the contraction response easy to observe and quantify (**Figure 1B**). When *Stentor* cells are exposed to a repeated stimulus, they will gradually stop responding to it, a response that shows many of the features that characterize habituation in other organisms (Wood 1970). For example, if cells are habituated to a given stimulus, they will still respond to stronger stimuli, showing that the response is not simply exhaustion; and cells habituate more rapidly and extensively to weak or high frequency stimuli.

**Figure 1.**
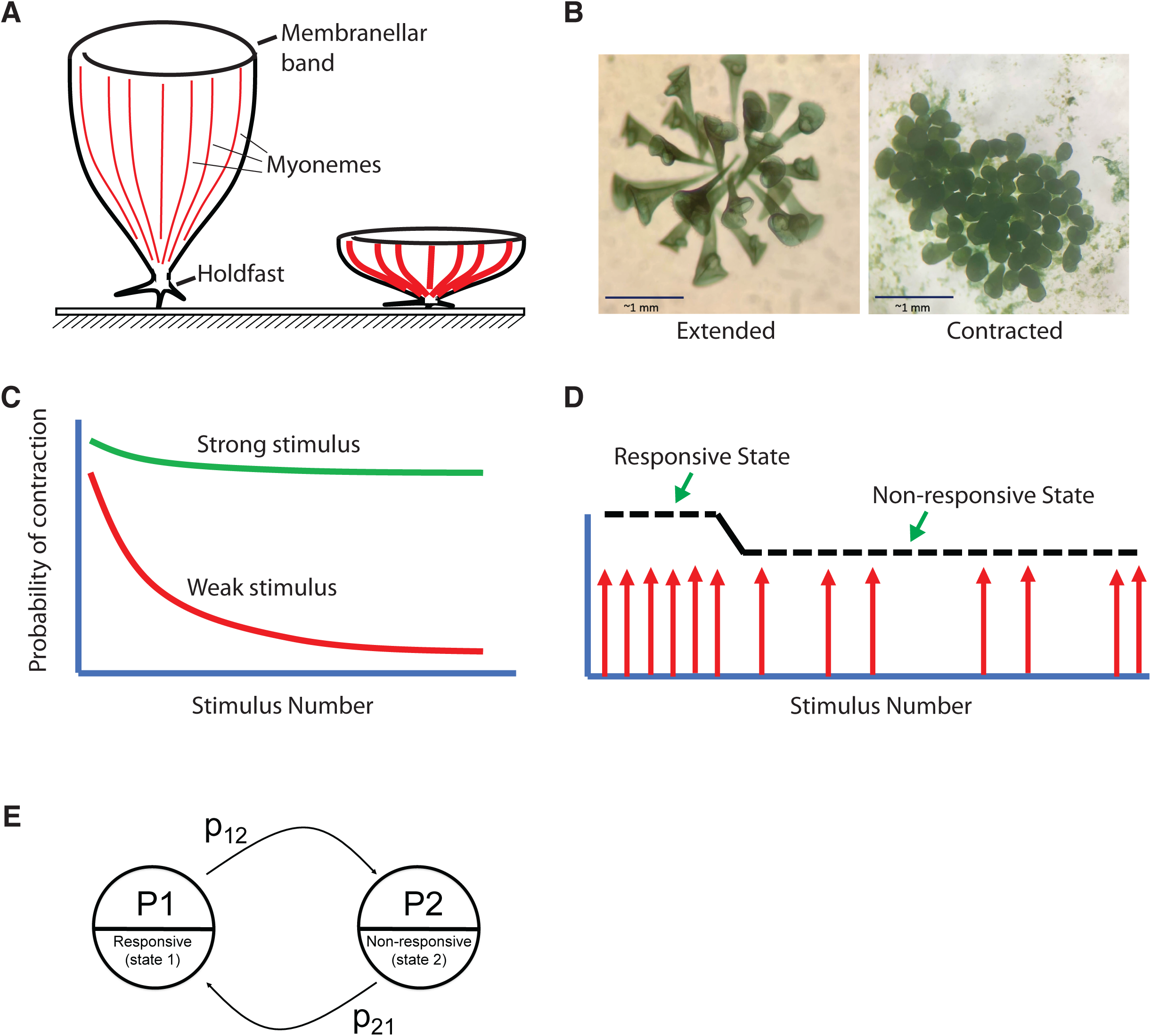
Habituation in the single-celled organism *Stentor coeruleus*. (**A**) When not disturbed, a *Stentor* cell extends into a cone shape, attached at its base to a leaf or other surface by means of a holdfast, and allowing it to draw in food using flow generated by a ring of cilia at the anterior end. When a *Stentor* cell experiences a mechanical stimulation, such as from a predator, it contracts by means of myonemes, which consist of long bundles of contractile EF hand proteins (shown in red). (**B**) Images of a cluster of *Stentor* cells in the extended and contracted states. (**C**) Habituation in S*tentor*: following repeated stimulation at a constant force, a population of cells gradually becomes less likely to contract in response to further stimulation. The rate and extent of habituation is less when cells are stimulated with larger magnitude force. (**D**) Habituation is step-like in individual cells. Red arrows indicate contraction events, showing that a cell typically starts out responding to every stimulus, but then appears to switch to a much lower rate of contraction. (**E**) A previously described two-state model (Rajan 2023a) suggested by the switch-like behavior of individual cells. The model represents a cell as being in two possible states, a responsive state which contracts in response to a stimulus with a probability P_1_, and a non-responsive state which contracts with probability P_2_. Each time a stimulus is applied, the cell can switch between the two states with probability p12 of going from the responsive to the nonresponsive state, and probability p21 of going from the nonresponsive back to the responsive state.

Compared to other cell types, *Stentor* is a convenient model system in which to study habituation at the single-cell level due to the ease with which the behavior (contraction) can be observed; the unambiguous nature of the essentially binary contraction response, which makes it easy to classify cells into contracted or non-contracted states with minimal uncertainty; the ability to monitor contraction and habituation in many cells at the same time in a single dish, which allows statistical numbers to be rapidly collected; and the short timescale necessary to observe habituation (20-30 min)., which allows experiments to be replicated with ease. The genome of *Stentor* has been sequenced (Slabodnick 2017) and RNAi methods are available to perturb gene expression (Slabodnick 2014). The molecular components of the *Stentor* cell have been cataloged by means of proteomics (Lin 2022) and transcriptomics (Sood 2022), all of which make it an excellent model system for exploring the mechanism of single-celled learning.

We previously confirmed the ability of *Stentor* cells to habituate, using a computer-controlled device to provide mechanical stimuli to *Stentor* cells (Rajan 2023a). Our results matched the results of Wood in that weaker stimuli led to faster and more extensive loss of the contraction response (**Figure 1C**). Unlike the work of Wood (1970a), in our experiments we tracked the responses of individual cells, and found that while a population of cells showed the gradual reduction in response probability that Wood had reported, when individual cells were observed, they appeared to undergo a more switch-like response from a highly responsive to a less responsive state (**Figure 1D**). By using model-fitting procedures including hidden Markov model reconstruction, we found that a two-state model (**Figure 1E**) could account for the observed behavior, but only if the reverse transition probability, from the non-response back to the responsive state, was extremely slow. The slow reverse transition rate was necessary to account for the fact that cells appear to take a single step to the non-response state and do not flip back and forth between the states during the training period.

Our previous study showed that a two-state model is sufficient to account for the learning behavior of single cells receiving stimuli at a constant frequency. This model can correctly match the dynamics of habituation in populations of cells, the step-like response observed from individual cells, and allowed the effect of force on habituation rate to be represented as an input that varies one of the transition rates (Rajan 2023a). It is worth noting that this last point, the ability to account for the force dependence of habituation rate, was something that had to be directly built into the model parameters by allowing one rate (the forward transition rate p12 from the responsive to the non-responsive state) to vary as a function of the force. This was not a prediction that followed automatically from any underlying model, just an assumption that needed to be added to allow the model to fit habituation data at both force levels. The force dependence of habituation was therefore something that the two-state model could accommodate, but only by adding an additional assumption.

Further experiments showed that the two-state model with force-dependent forward transition is NOT sufficient to represent the response of cells to different stimulus periods, nor could it represent the response recovery after cessation of the stimulus. With respect to different stimulus periods, it was found that when stimuli were applied at a very high frequency (1 per 1.2 seconds rather than 1 per minute), the habituation rate was higher when expressed in terms of time, but lower when expressed in terms of stimulus number. This is not consistent with a two-state model that has the chance to switch states with each stimulus, and suggested that the system was responding not just to the stimulus sequence as a discrete sequence of inputs, but also to the timing with which they were received.

With respect to response recovery, it is known that once the habituating stimulus ceases, *Stentor* cells will gradually return to the original pre-habituation response probability with a half-life of tens of minutes (Wood 1970a). In the two-state model, response recovery is driven by the reverse transition (probability p21 in **Figure 1E**) from non-responsive back to responsive state. When this parameter was adjusted to allow a good fit of single cell habituation data, the best fit value was extremely low (0.001 min^-1^) which was consistent with the fact that virtually all cells were observed to undergo a single transition to the non-responsive state, and never back to the responsive state. But using such a low value of the reverse transition probability means that the response recovery should take place with a half-life of 5 hours, rather than the 20 minute halftime observed in the same experimental measurements used to fit the transition rates (Rajan 2023a). This disparity between the reverse transition rate needed to explain the habituation curves during learning, and the transition rate needed to explain the response recovery during forgetting, was apparently a serious flaw in terms of the model’s ability to account for experimental data.

Thus, the two-state model has (at least) three major shortcoming when faced with actual data: (1) it does not predict force dependence, but instead requires this to be added as an assumption; (2) it is not compatible with the dependence of habituation rate on stimulus frequency; and (3) it does not allow a realistic response recovery rate when its parameters are chosen so as to represent single-cell habituation dynamics.

Another issue with the two-state model is implementation - although a model with two states is in some sense the simplest possible model, this view of simplicity comes more from the perspective of theoretical computer science, in which the finite state machine forms the lowest rung of a hierarchy of increasingly complex computational models, and the two-state model is the simplest non-trivial finite state machine. While we believe that there may be many contexts in which such a theoretical computer science perspective is appropriate for understanding cell behavior (Condon 2018), there is always the question of how to generate such states at a molecular level. In fact, implementing a two-state with molecules is non-trivial. Certainly, there are a large number of examples in which a single molecule can switch between two distinct conformational states, and such molecules play important roles in a huge number of biological processes (Philips 2020). But for a molecular switch to create a two-state system at the level of a whole cell would require a way to coordinate the activity of many molecules so that they all switch into one state or another in a synchronous fashion, which would require additional molecular machinery. Thus, a two-state switch-like behavior at the level of an entire cell may not actually be that simple to implement.

From a molecular perspective, a much simpler model would actually be one in which a population of molecules switch state independently, leading to a continuous rather than switch-like response. For example, we can envision a mechanism in which the mechanoreceptor itself, or some downstream molecule involved in signal transduction, is degraded or inactivated with a probability that depends on the current stimulus level, such that continued stimulation would lead to a gradual loss of receptor function. Here we investigate such a model in which we represent the inactivation and degradation of a molecule involved in mechanical force transduction, coupled with a thresholding mechanism based on previously published electrophysiology studies (Wood 1970b; Wood 1988a; Wood 1988b), and show that indeed such a mechanism can produce a switch-like behavioral transition in the cells even though the underlying "learning" process is based on a continuous variable representing, in this case, the quantity of functional mechanoreceptor located on the cell surface. In this sense it is just as good as the two-state Markov model. Unlike the two-state model, the receptor inactivation model directly predicts, without any additional assumptions and using a single parameter set, the dependence of habituation rate on force, the effects of increased stimulus frequency, and the timescale of response recovery. We further test this model using new experiments specifically designed to test model predictions, including an evaluation of “rate sensitivity” in terms of a dependence of response recovery rate on stimulus frequency. Unlike many other habituating organisms, *Stentor coeruleus,* under the conditions we tested, does not appear to display rate sensitivity. Based on these results, we suggest that the receptor inactivation model, while still highly simplistic, is not only more biologically realistic, it is also more effective as a predictive model, compared to our previous two-state model.

## Results

### Habituation and response recovery in a receptor-inactivation model

As a first step towards modeling possible biochemical pathways that might produce habituation, we started with a highly simplified scenario (**Figure 2A**) in which a mechanoreceptor linked to an ion channel responds to mechanical stimulation to open a channel and allow ions to flow across the membrane, thereby changing membrane potential. If the membrane potential crosses a threshold, calcium channels open allowing calcium to enter the cell. It is known that the contraction of *Stentor* is driven by myonemes composed of calcium-binding EF hand proteins (Huang and Pitelka 1973; Maloney 2005). For the purposes of this model, we assume that the calcium channel opening is an all or nothing response, such that whenever the membrane potential crosses a threshold, a contraction response is recorded. This assumption is consistent with prior demonstrations of a classical action potential in *Stentor* (Wood 1970b). We do not, however, explicitly model the detailed molecular components that generate the action potential, instead we just make the assumption that the action potential will be triggered, and cause cell contraction, whenever the voltage crosses a threshold.

**Figure 2.**
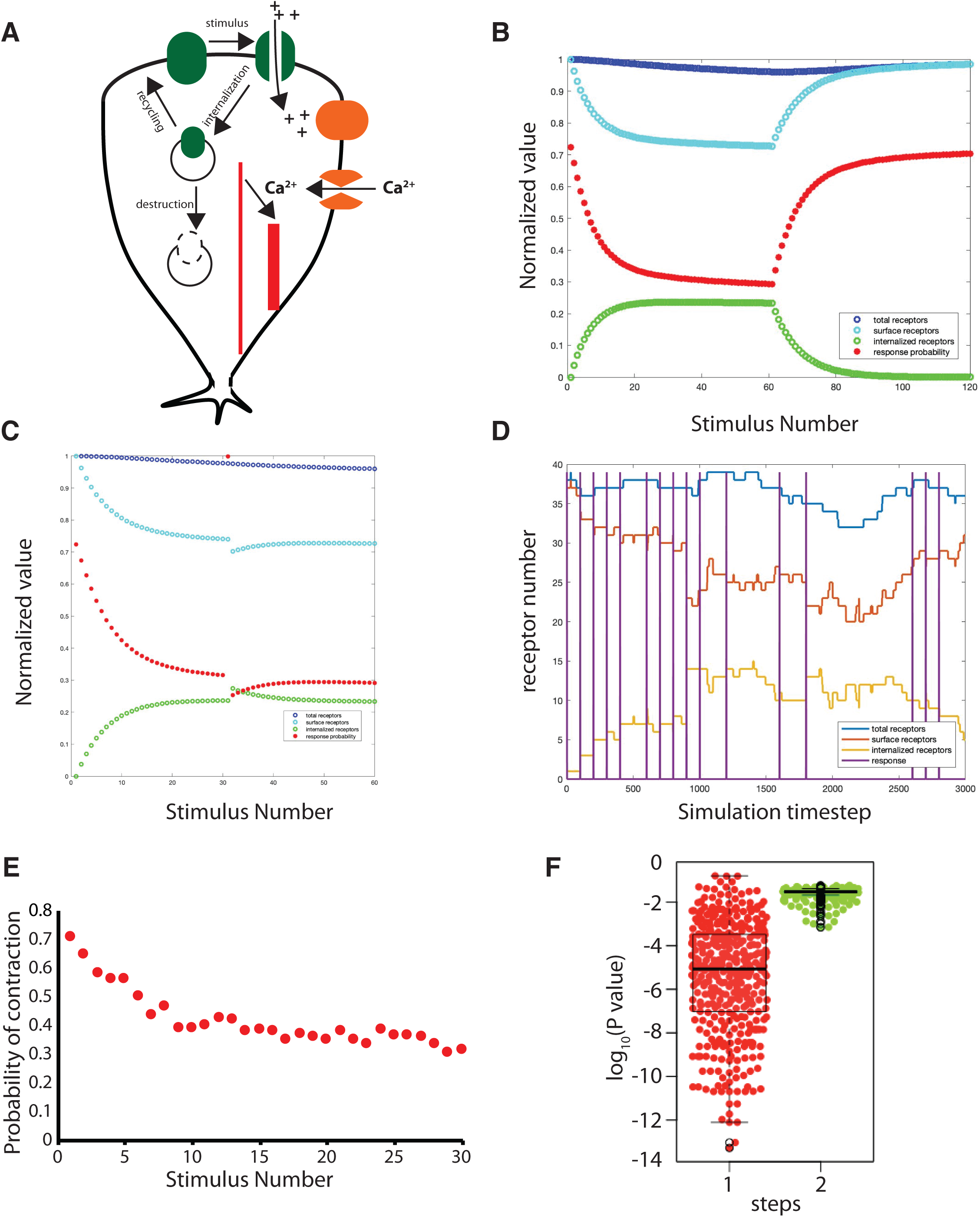
Biochemical model for habituation in *Stentor* based on activity-dependent receptor inactivation. (**A**) Diagram of model showing mechanoreceptors at the cell surface modeled as ion channels fluctuating between open and closed states. While in the open state, the receptor/channel can be internalized by endocytosis. Internalized receptors are subject to degradation, but can also be recycled back to the plasma membrane. Ion flow through the channels alters the membrane potential, which can trigger opening of a voltage dependent calcium channel. Influx of calcium leads to cell contraction. (**B**) Simulation of deterministic model for *Stentor* habituation. (blue) total receptor number, normalized to maximum value. (green) number of internalized receptors, normalized to maximum total receptor number. These receptors are the ones that are subject to degradation. (light blue) number of receptors on surface, normalized to maximum total receptor number. These receptors are the ones that can response to a stimulus. (red) probability of contraction. This simulation modeled a series of 60 stimuli applied at a frequency of 1 per minute. The reduced probability of contraction with successive stimuli indicates habituation. After 60 stimuli, the stimulus ceased and the probability of contracting if a stimulus were to be applied was then calculated for 60 more intervals, to quantify the response recovery. Return of the contraction probability to its initial value indicates recovery of response (forgetting). In this simulation, both habituation and response recovery occur on a timescale comparable to what has been reported in actual cells. (**C**) Lack of dishabituation in the receptor inactivation model. In this simulation, using the same parameters as panel B, 20 stimuli were applied to achieve habituation, followed by a single stimulus ten times greater, which caused a close to 100% contraction probability. After the single strong impulse, 20 more normal force stimuli were applied, and the response probability immediately shifted back to its previous habituated level, indicating that the model does not show dishabituation or sensitization. **(D)**. Example result of a stochastic simulation of receptor inactivation model. (blue) total receptor number, (orange) number of internalized receptors, (red) number of surface receptors, (purple) contraction events. In this simulation, stimuli were applied at a frequency of one per minute, and the simulation timestep was 0.01 minutes. Note that receptors can become internalized every time a stimulus is applied, regardless of whether the cell contracts, which is why the orange curve sometimes fluctuates upwards in between contractions. (**E**) Habituation in the stochastic implementation of the receptor inactivation model. Plot shows response probability averaged over 400 simulations confirming that habituation similar to that seen in actual cells occurs in the stochastic version of the model using the same parameters as panel B. (**F**) Statistical test for switch-like behavior in the stochastic model. Beeswarm plot shows p values for test for one and for two steps, indicating that a majority of cells show strong support for a single step-like response.

We further incorporate the fact that many receptors undergo inactivation and degradation following activation, in some cases via endocytic internalization and in other cases by proteolytic degradation (Kholodenko 2003, Shankaran 2007, Lin and Man 2013, Rosendale 2017). A wide variety of receptors and channels are known to be able to mediate mechanosensation in different organisms and tissues, including NOMPC/TRP channels, DEG/ENaC channels, and Piezo channels (reviewed in Katta 2015). We do not know the molecular identify of the *Stentor* mechanoreceptor, but among the various classes of mechanoreceptors in other species, many are part of receptor and channel families known to undergo internalization, ubiquitination, and other post-translational modifications following activation (reviewed in Estadella 2020). In our model, we focus on endocytic internalization, but alternative models could be developed for other mechanisms, that would be essentially similar.

We model receptor internalization by representing two pools of receptor, one located at the cell surface and one internalized, with only the former able to contribute to mechanosensation or changing the membrane potential. Internalized receptors can be either recycled to the plasma membrane, or destroyed, each with its own rate. Both surface and internalized receptors also undergo basal degradation at the same rate, which is lower than the internalization-mediated destruction rate, and the total receptor number increases over time due to receptor synthesis, which is assumed to occur at a constant rate and adds to the surface receptor pool. Details of the model, including the model parameters that describe receptor protein dynamics, input-output function for receptor activation, and electrical coupling, are given in Methods. We emphasize that the model parameters used in our simulations are not based on actual measurements in *Stentor* (or in fact any other organism), nor is the model intended to accurately reflect biochemical details in actual cells. Rather, the model is intended to test to what extent such a simple picture of the known molecular players involved in the contraction process may, or may not, be able to explain known features of *Stentor* habituation.

Simulations of this model show that indeed habituation can be accounted for by this simple mechanism (**Figure 2B**). As plotted in the graph, during continuous stimulation, the number of surface receptors (light blue) gradually decreases, accompanied by an increase in internalized receptors (green), leading to a correlated decrease in response probability (red). After the stimulus ceases (after 60 stimuli), the response probability gradually returns back to its initial value, driven by the synthesis of new receptors to replace those that had been lost by degradation. The parameter choice used in this simulation yielded an initial response probability, habituated response probability, habituation timescale, and timescale of response recovery (forgetting) that are comparable to what has previously been observed experimentally (Wood 1970a; Rajan 2023a).

The ability to account for the response recovery timescale is notable. One major difficulty with the two-state model that we previously proposed is its inability to fit both the habituation rate as well as the response recovery rate that are seen in actual *Stentor* cells (Rajan 2023a).

The reason is that in order to account for the switch-like behavior seen in single *Stentor* cells, it is necessary for the reverse transition rate (from non-responsive to responsive) in the two-state model to be extremely small. But this same rate constant is what determines the response recovery after cessation of stimuli in that model. In contrast, the receptor-inactivation model accommodates the observed time-scale of response recovery after stimuli are stopped (see **Figure 2B** timepoints 31-60).

One unusual feature of habituation in *Stentor*, that is distinct from habituation in most animal species, is the lack of dishabituation by strong stimuli, i.e. the ability of a single strong stimulus, when applied to a habituated organism, to restore responsiveness to weaker stimuli to which it had become habituated. This is a common feature in many cases of habituation (Thompson and Spencer 1966), but experiments by Wood have shown that it does not occur in *Stentor coeruleus* (Wood 1970a). In the receptor inactivation model, there is no mechanism for a strong stimulus to relieve the habituated state. If anything, applying a strong stimulus, which tends to drive internalization of more receptors, would only lead to stronger habituation. Simulations of the model confirm this intuition. As seen in **Figure 2C**, when a single strong stimulus is applied after a series of low for habituating pulses, it drives a contractile response because the stimulus is strong enough to activate a large enough number of receptors to cross the activation threshold, but after that pulse, when the stimulus returns to the previous lower magnitude, the response drops back down to what it would have been if the strong stimulus were never applied. Thus, our model is in agreement with the known lack of dishabituation by strong stimuli in *Stentor*.

### Receptor inactivation model recapitulates step-like habituation response of individual cells

One of the main experimental facts that led to the two-state model was the observation that even though a population of cells shows a graded response, individual cells showed a step-like response, apparently switching from one state in which the response probability was high, to a new state in which the response probability was low. The receptor inactivation model might seem inconsistent with this observation, since it is based on a graded internal variable in the form of the surface associated receptor number. But it is important to note that this graded internal variable is then read out by a non-linear process involving a thresholding operation. As the graded variable drops below a threshold, a sudden drop in response may be observed, which we hypothesized could cause the system to display a switch-like behavior despite the underlying continuous state variable. To test this idea, we carried out a stochastic simulation of the receptor inactivation model (see Methods). As shown in **Figure 2D**, this model incorporates fluctuations in the rate of receptor degradation and synthesis, and can simulate the response of individual cells as shown by the spiking response indicated by the purple bars. It can be seen in the example shown in **Figure 2D** that a cell that initially contracts on every stimulus appears to switch to a lower frequency of response. When this simulation is repeated many times, the average response of the simulations recapitulates the graded response of the population (**Figure 2E**). We previously described a statistical test for stepping (Rajan 2023a) based on the use of Fisher’s exact test to first infer the most likely time point at which a step might occur and then test the null hypothesis that the frequency of response is the same before and after the inferred step. Low P-values in this test indicate that the frequencies are unlikely to be the same and thus that a step has occurred. We previously applied this test to experimental data on *Stentor* habituation and found that there was strong statistical support for most cells taking a single step but not two or more steps (Rajan 2023a). Stochastic simulations of the receptor inactivation model showed the same result, with most cells appearing to take a single step based on the same statistical test which showed strong support for one step but little support for two or more steps (**Figure 2F**). Thus, the biochemical receptor inactivation model is capable of accounting for not only the gradual reduction in response in a population of cells, but also the apparent step-like response of single cells during *Stentor* habituation.

### Receptor inactivation model recapitulates dependence on magnitude of stimulus

One highly conserved aspect of habituation that is also seen in *Stentor* is the fact that habituation takes place more rapidly and more extensively in response to low force stimuli compared to higher force stimuli, such that with sufficiently strong stimuli, habituation barely takes place at all on the timescale of an hour (Wood 1970a). In our previous two-state model (Rajan 2023a), this force-dependent difference in habituation rates had to be accommodated by assuming that one or more parameters in the two-state model were inherently force-sensitive. Parameter fitting suggested that the main parameter affected by the force level was the forward transition rate (from the responsive to the non-responsive state).

When we simulated the receptor-inactivation model, we found that in fact a single set of parameters could account for the habituation dynamics of two different force levels, such that low force stimuli caused extensive habituation (**Figure 3A**) while high force stimuli caused little habituation (**Figure 3B**), a trend that extended over a wide range of stimulus magnitudes (**Figure 3C**), thus recapitulating what has been seen in experiments, without the need to arbitrarily retune parameters to model different levels of force as we had previously done for the two-state model (Rajan 2023a). Thus, the receptor inactivation model can explain this phenomenon in a more parsimonious way compared to our previous two-state model.

**Figure 3.**
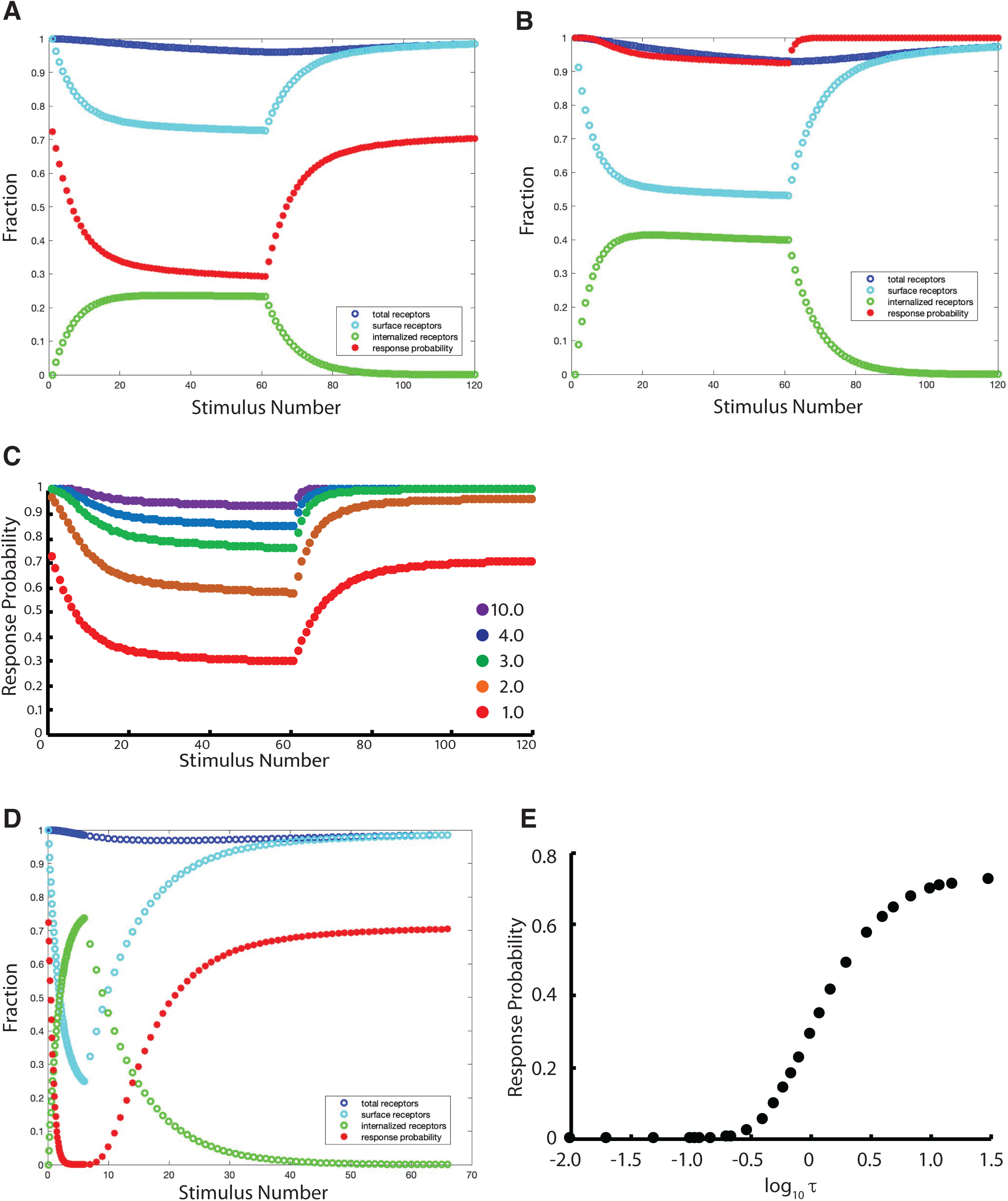
Predicted effect of stimulus magnitude and frequency on habituation. (**A,B**) Simulations of two different force stimuli with the stimulus in panel B tenfold higher (10 force units) than that in panel A (1 force unit). With the high force stimulus, receptor number still decrease (blue curve) but this does not lead to a large drop in response probability because the large force is able to trigger a larger fraction of the remaining receptors. (**C**) Response curves for different stimulus forces, showing that habituation is faster and more extensive for smaller forces, and slower and less extensive for larger forces. (**D**) Simulation using the same stimulus magnitude as panel A, but with a ten-fold higher frequency. Comparing panels A and D it is apparent that the rate and extent of habituation is greatly increased with this high frequency stimulus. (**E**) Response probability at the end of the stimulus sequence versus the inter-stimulus interval (τ) of the stimulus sequence. In all case, the same total number of stimuli were applied.

### Dependence of habituation on frequency of stimulation

As discussed above, we previously showed experimentally (Rajan 2023a) that stimuli applied at a higher frequency (1 per second versus 1 per minute) led to faster habituation in terms of the actual time, but slower habituation in terms of the number of stimuli - that is, habituation to the same extent required a larger number of stimuli at high frequency, but because the time between stimuli was so low, the overall time for habituation was less than seen with lower frequency stimulation. This phenomenon was not explainable in terms of the two-state model. In contrast, as shown in **Figure 3D**, our simulation predicts faster habituation for stimuli applied at lower frequencies (compare to **Figure 3A** which uses the same force magnitude but lower frequency).

Wood (1970a) reported that compared to the standard stimulus period of 1 per minute, stimuli of lower frequency produced less of a decrease in response probability at steady state. Our simulation results recapitulate this result (**Figure 3E**). Conversely, we also found that higher frequency stimulation could produce more extensive habituation in the sense that the response probability at steady state was lower than with low frequency stimulation.

Taken together, our simulations indicate that the highly simplistic biochemical model for habituation based on receptor inactivation can account for many of the previously reported phenomenological features of habituation in *Stentor*, namely: the habituation response for a population of cells; the consistent time-scales of both habituation and response recovery; the lack of dishabituation; the apparently step-like response of individual cells, faster and more extensive habituation in response to weaker stimuli compared to stronger stimuli; and the dependence of habituation extent on stimulus frequency. Many of these features could not be accounted for by the previous two-state model. In light of the apparently reasonable predictions of the receptor inactivation model, we next used the model to design new experiments to further probe the nature of habituation in *Stentor*.

### Experimental test of model prediction: high force stimuli followed by low force stimuli

A comparison of **Figure 3A** and **Figure 3B** show an interesting feature, that although the cell fails to habituate significantly to a high force stimulus, the total receptor number still decreases during high force stimulation. The reason that habituation is not observed in this model is that even though the receptor number drops substantially, if the stimulus is strong enough, the remaining receptors become fully activated and keep the system above the threshold for triggering an action potential. The fact that receptor number drops during exposure to high force stimuli that do not yield observable habituation leads to a prediction: if we were to follow a series of high force stimuli with a series of low force stimuli, the cell should show a reduced response to the first low force stimulus sequence compared to if it were applied to a naive cell. **Figure 4A** illustrates the experiment, and **Figure 4B** shows the result of simulation. It can be seen that the model predicts a reduced response to the first low-force stimulus following the high force stimuli.

**Figure 4.**
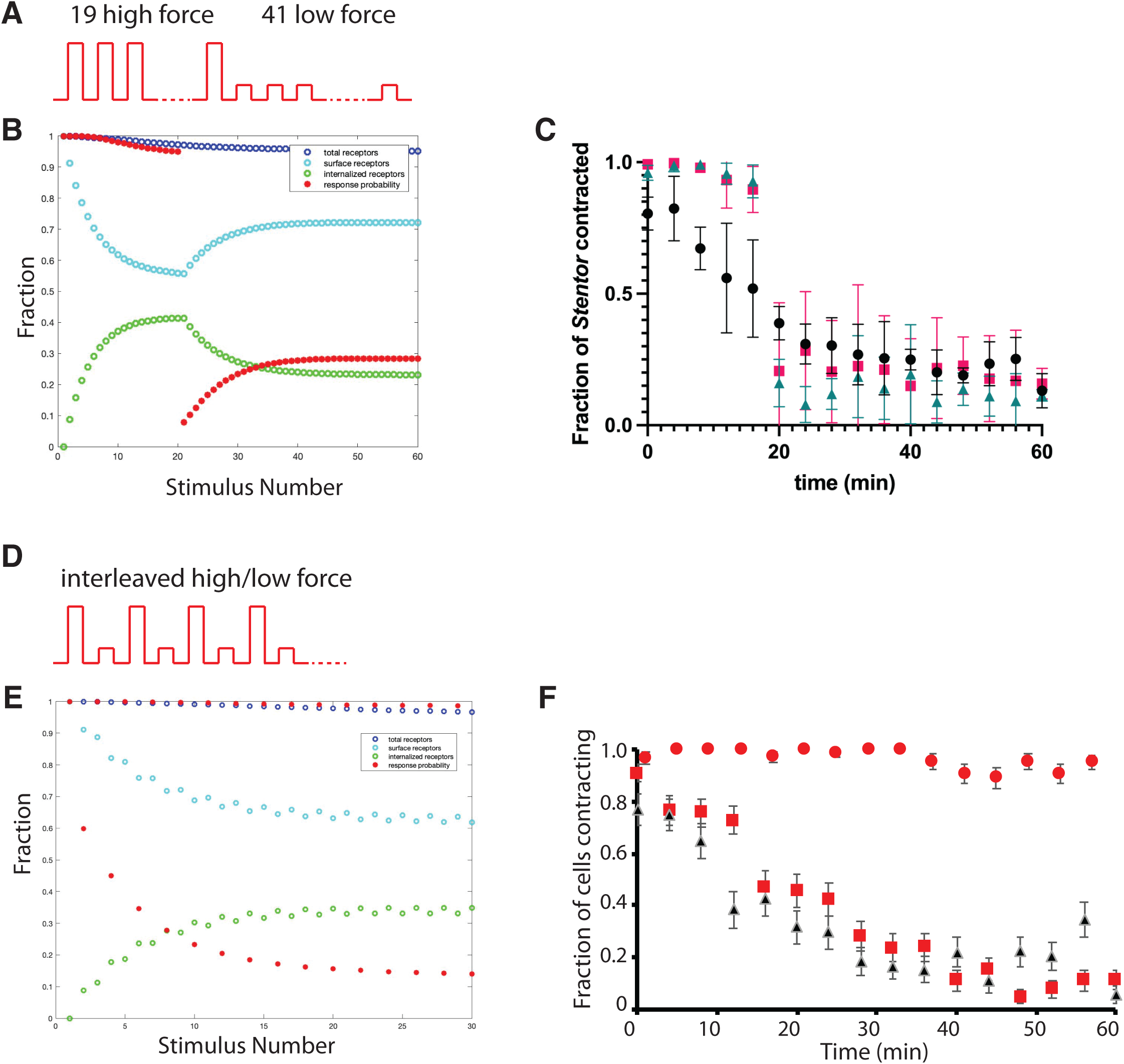
Habituation of low-force response during high-force stimulation: simulations and experiments. (**A**) Experimental design to test model prediction that a cell exposed to high force stimuli should show reduced response to subsequent low force stimuli. Experiment involves presenting series of 20 high force stimuli, followed by 40 low force stimuli. (**B**) simulation showing that a series of high-magnitude stimuli that cause little observable habituation to the high force stimulus lead to a decreased response to a lower force stimulus. Light blue markers show that receptor number on the cell surface drops during the high force stimulus part of the sequence, which explains why the response to the subsequent low force stimuli starts out at a low level. (**C**) Experimental measurement based on the experimental design of panel A. In this experiment, a sequence of 19 high force taps (red squares; Force level 3; N=59-85) was applied at a rate of 1 tap per minute, after which 41 low force (force level 4) taps were applied at 1 tap per minute. Despite the fact that the cells had never experienced a low force tap, but only high force taps to which they do not significantly habituate, they indeed showed a reduced probability of response to the low force taps, comparable to what is seen in equivalent experiments carried out using only low force taps (black circles; force level 4, N=51-87). A similar effect is seen when an even higher force (level 2) is used for the first 19 taps followed by the same low force (level 4) for the remaining 41 taps (green triangles; N=70-84). Error bars signify standard error of the mean; each experiment was done in triplicate. N values indicate the range for the number of cells anchored in the field of view over the course of each series of experiments. (**D**) Alternative experimental design to test whether high force stimuli can cause habituation of low force stimuli, using interleaved high and low force stimuli. (**E**) Simulation of interleaved high and low force stimuli, showing that the response to the low force stimulus undergoes habituation but does not cause habituation for the large force stimulus. Light blue markers show that surface receptor number drops after both types of stimuli. (**F**) Experimental measurement of interleaved high (red squares; force level 3) and low (red circles; force level 4) magnitude stimuli (number of cells analyzed N=59-72 across the experiment) applied at an overall rate of one stimulus per minute, with the high force taps alternating with the low force taps. Black triangles indicate a control experiment using only the lower force level taps with identical timing (force level 4; N=51-60). Error bars signify standard error calculated from binomial statistics. As predicted by the model, the response to the low force stimuli shows extensive habituation even though the response to the alternating high force stimuli does not.

To test this prediction, we used our stepper motor-based habituation device (Rajan 2023b), which allows the stimulus magnitude to be varied during the course of an experiment, to stimulate cells with a high force stimulus for 19 iterations, followed by 41 low force stimuli. The results are plotted in **Figure 4C**. Also plotted, for comparison is the result of a comparable experiment but using only low force stimuli. It can be seen that despite little reduction in response during the high force stimulus, the response to the subsequent low force stimulus is indeed reduced, to a level comparable to what would have been observed if low force stimulation had been used throughout the experiment. A similar effect was seen using an even higher force for the initial training. This experimental result confirms the prediction of our receptor inactivation model. This result is fundamentally incompatible with our previous two-state model, in which the low transition probability to the non-responsive state during high force stimulation would mean that when the low force stimuli were applied, the response rate would still be high initially because most cells would still be in the responsive state, and would only start to decrease after successive low force stimuli were applied.

### Experimental test of model prediction: interleaved high and low force stimuli

The previous simulation and experimental comparison supports the idea that there is a hidden variable, which in the model represents the quantity of receptors, which can change even in response to stimuli that have little observable effect on habituation. As an alternative way to compare the model with this prediction, we simulated an experiment in which strong and weak stimuli were alternatively applied in an interleaved fashion (**Figure 4D**). As shown in **Figure 4E**, simulations of such an interleaved weak/strong stimulus experiment predict that the response probability to the low magnitude stimulus should still show habituation but this will not carry over to the strong force stimulus. We note that this prediction is entirely different from the prediction made by the two-state model, in which once a cell is in the non-responsive state, it would show reduced response to both stimuli, thus the low magnitude stimuli would tend to drive faster habituation of the higher magnitude stimuli. We thus have a distinct set of predictions for the two models - the receptor inactivation model predicts that interleaved stimuli would show habituation of the responses to the low force stimuli, but not to the high force stimuli, whereas the two-state model predicts that interleaved stimuli would lead to faster habituation of the high force stimuli than would be seen if the high force stimuli were presented exclusively.

We tested the predictions of the two models experimentally in *Stentor* cells, by using our stepper motor habituation device to provide alternating low and high force taps. As shown in **Figure 4F**, the results match the prediction of the receptor-inactivation model, in that the response probability to the low magnitude stimuli showed a continuous decrease, while the response to the high magnitude stimuli continued to stay high comparable to what has been seen in high force only experiments.

### Experimental test of model prediction: subliminal accumulation

A common hallmark of habituation is subliminal accumulation, a phenomenon by which memory retention improves when training continues even after habituation has reached asymptotic levels (Thompson and Spencer 1966; Rankin 2009). The receptor internalization/degradation model proposed here involves two key processes that are likely to act on two different timescales: receptor internalization/recycling, and receptor protein degradation/synthesis. Simulations in which a stimulus is applied for a prolonged period, even after the point at which the cell has ceased to response (**Figure 5A**), show that the total receptor number continues to decrease during the prolonged training period, such that complete response recovery is delayed following cessation of stimulus, compared to simulations in which the stimulus is removed when the response drops to zero (**Figure 3D**).

**Figure 5.**
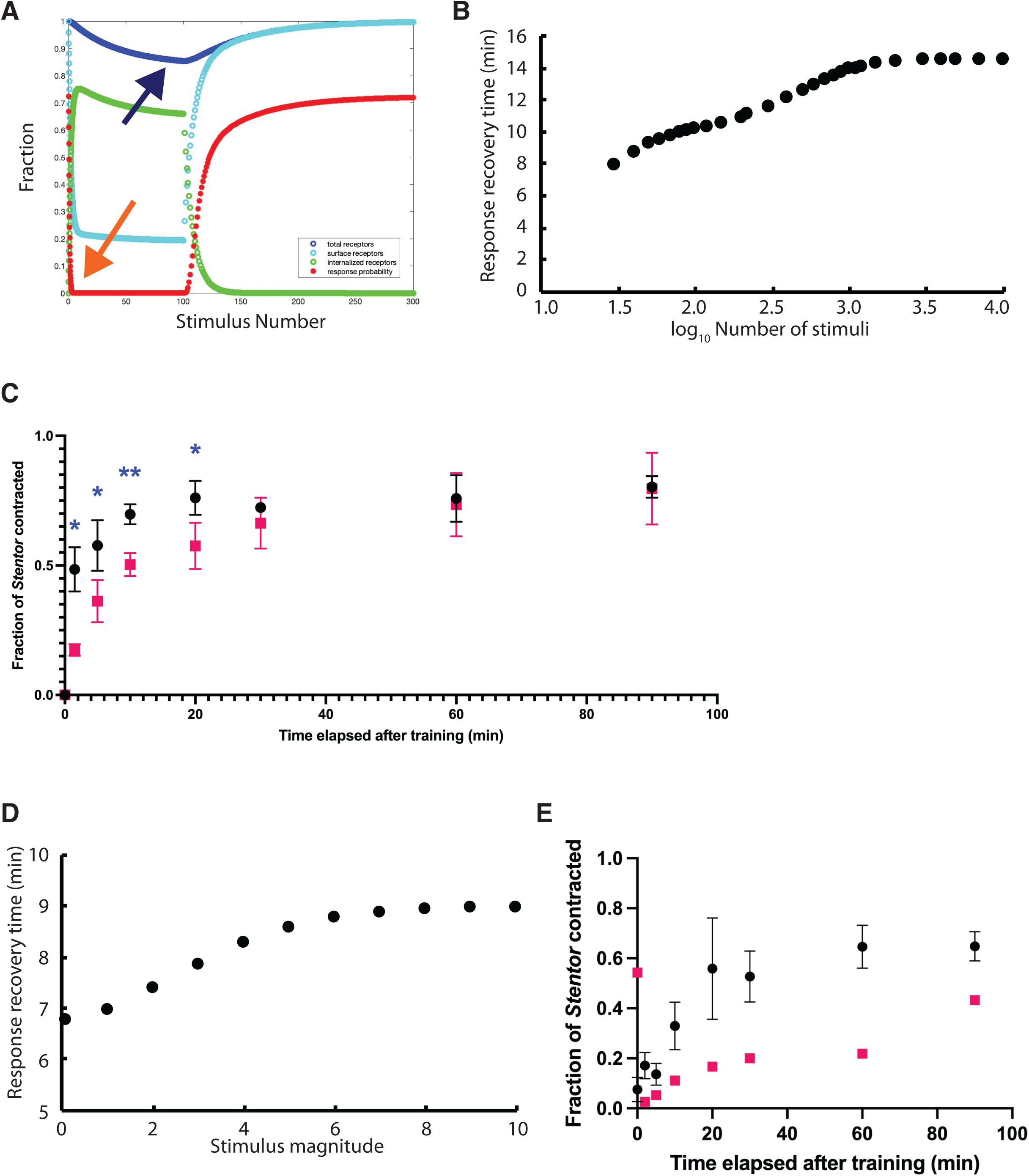
Dependence of recovery time on training duration (subliminal accumulation) and stimulus strength: simulations and experiments. (**A**) Simulation of extended habituation using high frequency stimulation to rapidly habituate cells to zero response (orange arrow), and then continuing the training beyond that point. During this extended training, the total receptor number begins to drop (blue arrow). The results is that the time taken to recover full response is substantially longer than in previous simulations that did not include the extended habituation period (compare Figure 3D). Figure indicates that for prolonged learning, the response recovery parallels the recovery of total receptor number by protein synthesis, in contrast to shorter duration training such as in Figure 3, where response recovery parallels recycling of receptor to the plasma membrane. (**B**) Simulation result plotting response recovery half-time versus the number of stimuli (force magnitude 1, period 0.1 min). Recovery time is defined here as the time for the response probability to recover halfway between the probabilities seen in fully habituated and fully naive cells. (**C**) Experimental measurement of response recovery following stimuli applied at a period of 1.2 sec for 12 hours (pink; N=35-44 across three experiments) versus 20 minutes (black; N=47-64 across three experiments). Error bars signify standard error of the mean; each experiment was done in triplicate. N values indicate the range for the number of cells anchored in the field of view over the course of each series of experiments. * p < 0.05, ** p < 0.01; Welch’s t-test. (**D**) Simulation results of response recovery half-time versus stimulus magnitude, for simulations using 1000 stimuli with a period of 1 min and a range of forces indicated. **(E)** Experimental measurement of response recovery following training with two different magnitude forces (black; N=21-45 across three experiments) force level 4 (low force), (pink) force level 3 (high force). Error bars signify standard error of the mean; low force experiments were done in triplicate. N values indicate the range for the number of cells anchored in the field of view during experiments.

This trend is confirmed by multiple simulations as plotted in **Figure 5B**. In order to test this prediction experimentally, we habituated cells with identical stimuli for either 20 min or 12 hours, after which cells in both training paradigms reached asymptotic levels of habituation, and then measured the response recovery (**Figure 5C**). The results match the prediction of the simulations in the sense that recovery takes longer for cells exposed to a longer duration of training.

### Experimental test of model prediction: memory retention following high force stimuli

Simulations also showed that increasing the magnitude of the training stimulus leads to a longer recovery time (**Figure 5D**), but the effect is less pronounced than the effect of training duration. Experimental measurements of recover following a series of strong versus weak stimuli (**Figure 5E**) showed a significant effect of stimulus magnitude on recovery time, which was much more significant than our simulations predict.

### Comparison of model predictions with experimental measurement of potentiation

The separation of timescales between internalization/recycling and degradation/synthesis that was evident in the simulations of Figure 5 suggested a possible basis for the phenomenon of potentiation, one of the commonly observed features of habituation in animals (Thompson and Spencer 1966), in which repeated rounds of training and recovery lead to an increase in habituation rate. Wood (1970) has reported such an effect in *Stentor*, although the extent of the effect was small. When we simulated repeated rounds of training separated by rounds of recovery (**Figure 6A**), we did notice that there was a slight drop in the initial response rate upon restoration of the stimuli after recovery. When habituation curves were compared for the first four training rounds, it was apparent that the first round of habituation showed an overall higher probability of response and a slower rate of habituation (**Figure 6B**), while the subsequent training rounds all showed similar response curves. We repeated the potentiation experiment ourselves (**Figure 6C**), and found results that were very similar to our simulations: the first round of habituation gave overall higher response probability and slower habituation, whereas the subsequent three training bouts were all similar to each other. The degree of potentiation seen in our simulations was not as strong as seen in our experiments, but we cannot rule out that some other parameter values might give a stronger potentiation effect.

**Figure 6.**
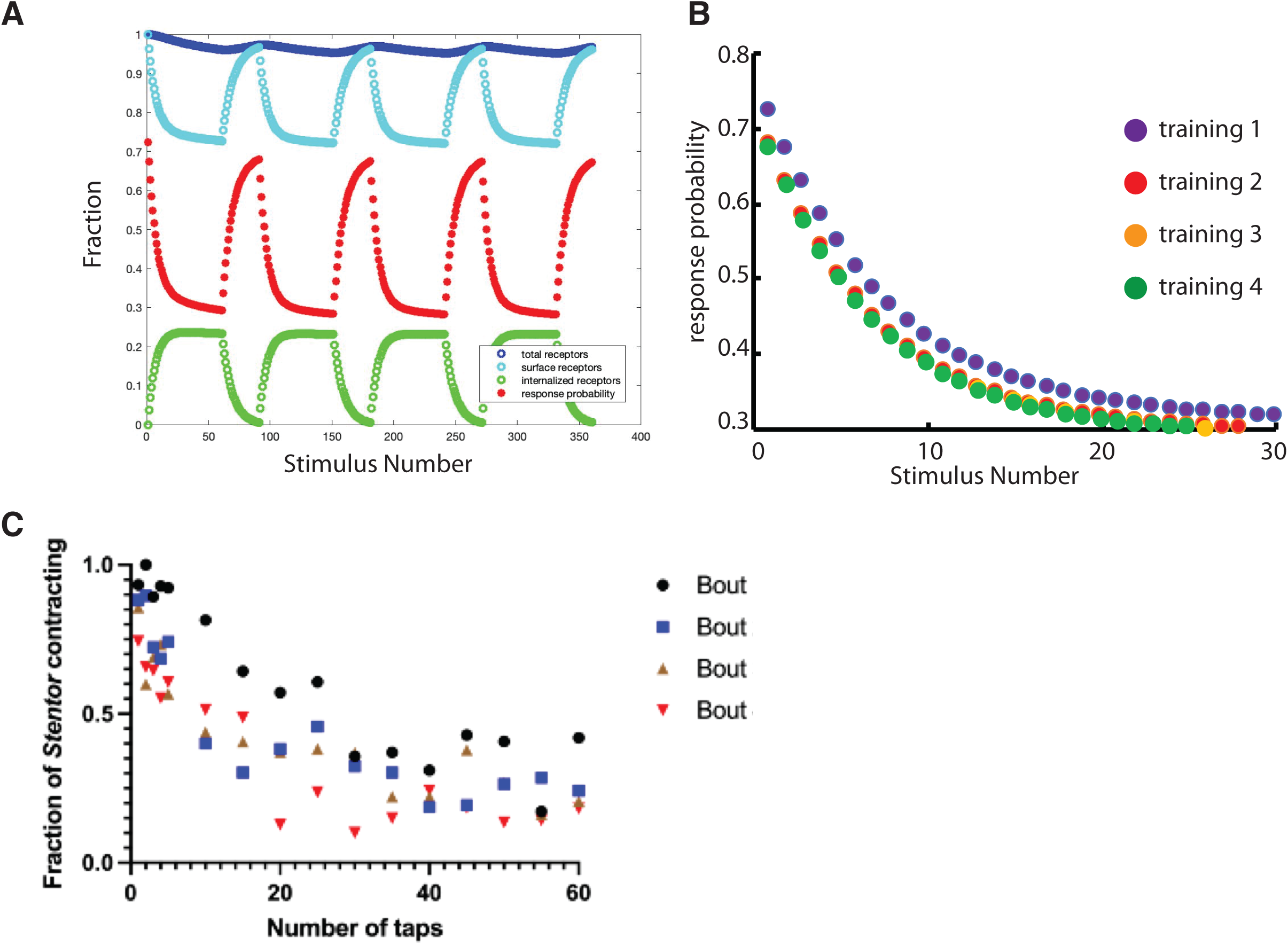
Faster habituation following cycles of habituation and recovery: simulations and experiments. (**A**) Simulation of repeated cycles of habituation (60 stimuli) followed by recovery (30 stimulus periods with no stimulus). (**B**) Response probability versus stimulus number for four successive bouts of training, showing just the first 30 stimuli. (**C**) Experimental test of habituation potentiation in which spaced repetition increases speed of learning. Each bout featured 60 minutes of training at a frequency of 1 tap/minute, followed by a 30-minute break: bout 1 (black circles; N=26-31), bout 2 (blue squares; N=29-35), bout 3 (brown triangles; N=29-37), bout 4 (red inverted triangles; N=31-40). N values indicate the range for the number of cells anchored in the field of view over the course of each training bout.

### Experimental test of model prediction: apparent lack of rate sensitivity on response recovery time

One commonly observed feature of habituation in animals is "rate sensitivity" which is defined as faster recovery of response following cessation of stimulus when the stimulus frequency was higher (Staddon 1993; Staddon and Higa 1996). Previous studies investigating the effect of stimulus frequency on habituation in *Stentor* (Wood 1970a; Rajan 2023a) reported effects of stimulus frequency on the rate and extent of habituation but did not address whether or not rate sensitivity occurs in this system.

Using our simulation of the receptor inactivation model, we calculated the predicted response recovery time as a function of the stimulus frequency. **Figure 7A,B, C** depict simulation results using the same parameter values and the same stimulus magnitude, but with progressively increasing stimulus frequencies. **Figure 7D** summarizes the response recovery half-times calculated for a range of stimulus frequencies. In contrast to **Figure 3E** which showed that the habituation extent depends on stimulus frequency, our model indicates that the response recovery time is changes very little as a function of stimulus frequency, and in fact shows a slight increase, rather than decrease, in recovery time for very high frequencies. The reason for this increase can be seen from the simulation results in **Figure 7A-C**, where the higher frequency stimulation leads to a more extensive receptor internalization, with a correspondingly greater drop in the response probability. Because higher frequency stimulation causes greater internalization of receptors in a given length of time, it takes longer to recover full response by recycling after the stimulation ceases. We note that this type of dependence, in which the response recovery time is greater for higher frequency stimuli, is the opposite of what has been reported for "rate sensitivity" in animal habituation. We conclude that the receptor inactivation model described here cannot account for the rate-sensitivity phenomena seen in other organisms.

**Figure 7.**
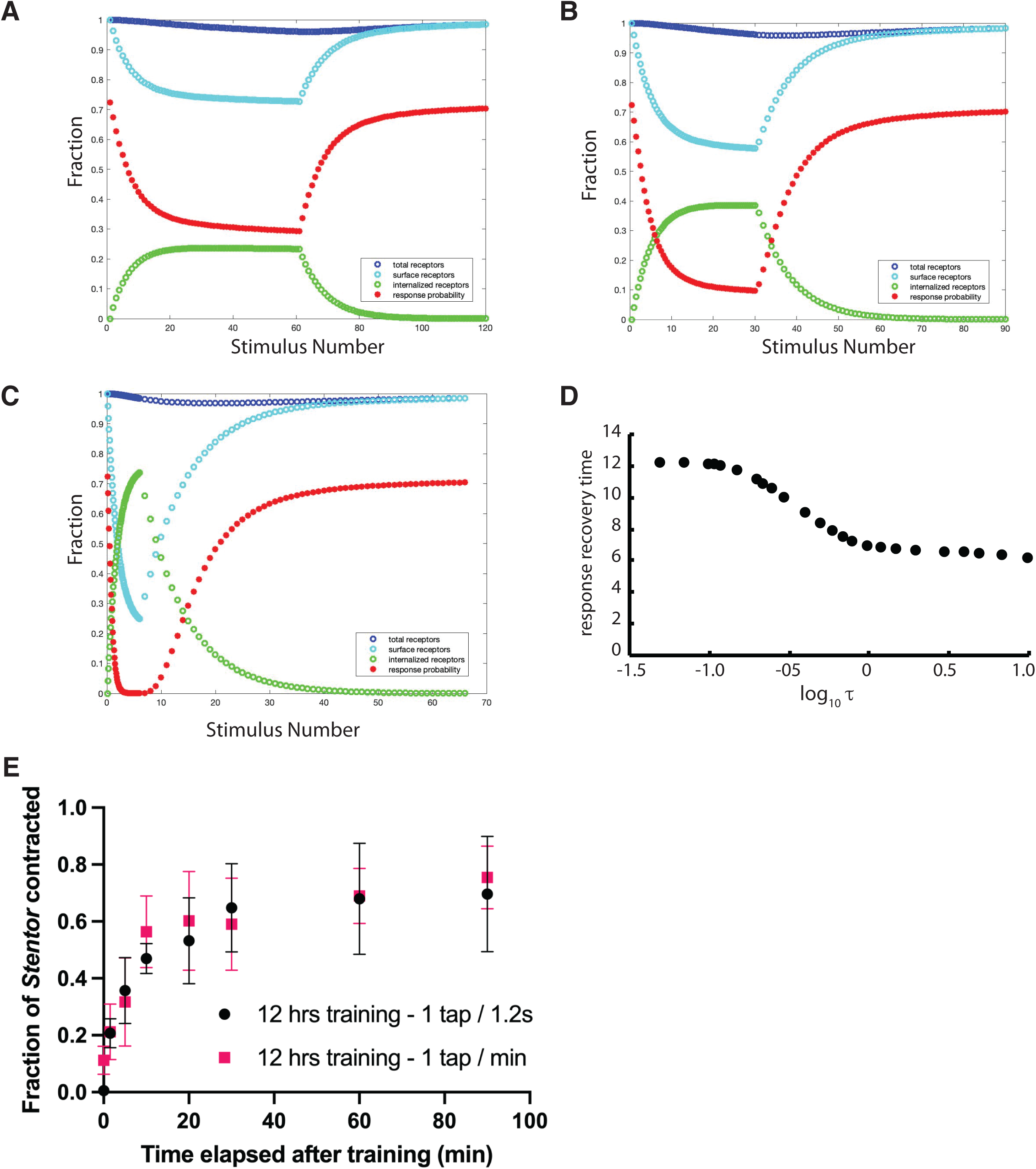
Stimulus frequency does not affect response recovery time: simulations and experiments. (**A,B,C**) Three simulations using the same model parameters and stimulus magnitude but different stimulus periodicities (A 1 minute, B 0.5 minutes, C 0.1 minutes). After the stimulus ends, the response recovery can be seen to occur at a similar timescale in the first two cases. With the extremely high frequency stimulation in panel C, response recovery takes longer than for the lower frequency stimuli. (**D**) Response recovery time versus stimulus frequency. For each simulation used to generate this curve, habituation was carried out until the response probability reached a steady state. (**E**) Experimental measurement of response recovery following habituation to two stimuli applied at frequencies of 1 tap per minute (red squares; N=25-47 across three experiments) or 1 tap per 1.2 seconds (blue circles; N=35-44 across three experiments), showing that the response recovery is virtually the same in both cases. Error bars signify standard error of the mean; each experiment was done in triplicate. P > 0.05 at each time point; Welch’s t-test.

We therefore asked whether the habituation of *Stentor* cells actually exhibits rate sensitivity, by habituating cells with the same force stimulus at two different frequencies: 1 tap per 1.2 sec versus 1 tap per minute. As shown in **Figure 7E**, the response recovery took place with virtually indistinguishable kinetics in both cases. These results suggest that *Stentor* may not display rate sensitivity, at least not under the parameters of our current experimental conditions, which would be consistent with the prediction of the receptor inactivation model.

## Discussion

### Evaluation of biochemical model with respect to empirical data about Stentor habituation

Habituation is defined as a decrement in response probability following repeated exposure to a constant stimulus (Harris 1943). However, over the years experimental psychology studies in various animals from invertebrates to humans have revealed a number of additional features that often accompany habituation. Thompson and Spencer (1966), in a comprehensive review of habituation in multiple animal model systems, defined nine widespread features that habituation displays. This list was updated in a more recent publication (Rankin 2009) which clarified the wording and concepts and added one more. Here we consider the extent to which these ten features can be explained by our model as well as the degree to which they are actually observed in experimental studies of *Stentor* habituation.

#### 1. Repeated application of a stimulus results in decreased response, usually as a decaying exponential function of the number of stimulus presentations

As shown in **Figure 2B**, our model shows the decrease in response, and the form is visually similar to an exponential, as explained by the fact that the mechanism for the response decrement is receptor internalization which occurs as a first-order process. Prior studies in *Stentor* also show such a decay curve during habituation (Wood 1970a; Rajan 2023a). So in this case, the model agrees with experiment, both of which agree with the summary of Thompson and Spencer.

#### 2. After the stimulus stops, the response recovers over time (forgetting)

As shown in **Figure 2B**, our model demonstrates response recovery when the stimulus ceases, taking place on a similar timescale has been documented in *Stentor* (Wood 1970a; Rajan 2023a). So again, in this case, the model agrees with experiment, both of which agree with the summary of Thompson and Spencer.

#### 3. As cycles of habituation and recovery are repeated, successive rounds of habituation take place more rapidly (potentiation)

Wood (1970) reported that after one round of habituation followed by a 6 hour rest period, when a second round of habituation was carried out, the habituation reached its minimum value in 40 minutes rather than 45 minutes for the first round, a small difference in habituation rate but one which was statistically significant. Our own experiments confirm that potentiation occurs in *Stentor* (**Figure 6C**). An effect of this type, which is a form of latent memory, requires that some molecular change persists on a time scale longer than the overt recovery of the response. In the receptor inactivation model, this longer time scale can in principle be provided by destruction of receptor protein, which then needs to be restored by protein synthesis, which is slower than the recycling process that dominates the response recovery after short periods of stimulation. As shown in **Figure 6A,B**, our simulations demonstrate a potentiation effect, which is manifested as a small difference between the habituation during the first round of training relative to successive rounds, all of which are essentially identical to each other. However, the extent is smaller than that seen in our experiments (**Figure 6C**). It is possible that some other choice of parameters might give a stronger effect, we have not explored this question since in this paper we have made every effort to use a single consistent set of parameters for all simulations.

#### 4. Higher frequency of stimulation leads to faster and/or more extensive reduction in response, and more rapid response recovery after the stimulus ceases

Our model recapitulates the first part of this effect (**Figure 3E**), namely the greater extent of habituation with high frequency stimulation, which has also been experimentally demonstrated in *Stentor* (Wood 1970a; Rajan 2023a). So in this case, the model agrees with experiment, both of which agree with the summary of Thompson and Spencer.

The second part of this feature, namely the increased rate of recovery when habituation is performed with higher frequency stimuli ("rate sensitivity") was not included in Thompson and Spencer’s list of nine characteristic features of habituation, but was added in their 2009 revision (Rankin 2009). Rate sensitivity has been observed in rats (Davis 1970), *Aplysia* (Byrne 1982), and *C. elegans* (Rankin and Broster 1992), but was not reported in Wood’s analysis of habituation in *Stentor* (1970). Our model does not show such behavior. On the contrary, the model predicts that higher frequency stimuli might lead to slightly slower, rather than faster, response recovery (**Figure 7D**). Our experiments presented here also failed to detect evidence for rate sensitivity in *Stentor* under the particular conditions we employed (**Figure 7E**), so in this case the model agrees with experiments in *Stentor*, but both disagree with reports on habituation in animals.

#### 5. Weaker stimuli produce faster or more extensive habituation compared to strong stimuli which, in some cases, may not yield habituation at all

Our model recapitulates this effect (**Figure 3C**), which has also been experimentally demonstrated in *Stentor* (Wood 1970a; Rajan 2023a). So in this case, the model agrees with experiment, both of which agree with the summary of Thompson and Spencer.

#### 6. Prolonged exposure to a stimulus after the point at which response has ceased leads to a slower response recovery

Our simulations of the receptor inactivation model clearly show this effect (**Figure 5B**), which is attributable to the two timescales of the model, such that the initial habituation is driven by receptor internalization, which takes place rapidly, but then as the stimulation duration is prolonged, receptor degradation occurs (**Figure 5A**), which then requires receptors to be synthesized de novo in order to achieve full response recovery. This effect was not reported by Wood (1970a), but our new experiments confirm that longer duration of training leads to a longer time for response recovery (**Figure 5C**). This is, therefore, another case where both the model, and experimental measurements in Stentor, agree with the criteria of Thompson and Spencer.

#### 7. Habituation to a given stimulus exhibits stimulus confers reduced response to other stimuli

Our model would not produce this effect given that the receptor that undergoes inactivation/degradation is presumably specific to the mechanical stimulus. Wood (1970a) has shown that the response to electrical pulses is not altered by habituation to mechanical stimulation, and that habituation to mechanical stimuli does not affect sensitivity to light stimuli and vice versa (Wood 1973), so in this case our model agrees with experiments in *Stentor*.

#### 8. Presentation of a strong stimulus can lead to recovery of response (dishabituation)

Lack of dishabituation to strong stimuli in *Stentor* has been previously reported (Wood 1970a). Our model also does not display dishabituation to strong stimuli (**Figure 2D**), so in this case the model agrees with experiments in *Stentor*, but neither agrees with the trend reported by Thompson and Spencer.

#### 9. When a strong stimulus that produces dishabituation is applied repeatedly, its effect diminishes

Because *Stentor* cells do not dishabituate to strong stimuli, this feature is not applicable.

#### 10. Some experimental protocols may produce habituation that persists for hours, days, or weeks (long term habituation)

In our model, the response recovery time is only a function of how small the number of receptors was at the end of the stimulus sequence, and the rate of receptor synthesis. There is no way to manipulate the stimulus sequence protocol to slow down the response recovery, other than driving the receptor number as low as possible. We therefore do not expect that the model as formulated could account for long term habituation. Long term habituation was not reported by Wood (1970a). We note that long term habituation is apparently only seen in a few special cases (Rankin 2009) and is not a universal feature of habituation in animals.

### Limitations and future directions for experimental tests of model predictions

Although we report here a lack of evidence for rate sensitivity in *Stentor*, we acknowledge the very real possibility that rate sensitivity might be a component of *Stentor* habituation under alternative experimental conditions and training regimes. Cells that undergo training for 12 hours at a frequency of 1 stimulus per minute (low-frequency) do not habituate as completely as cells that experience the same length of training at frequency of 1 stimulus per 1.2 seconds (high-frequency), such that the fraction of *Stentor* contracted at time zero post-training is slightly lower for the latter group (**Figure 7E**). Given that claims of rate sensitivity in other organisms are typically made when habituation has reached the same steady state level in both high-frequency and low-frequency training groups, this slight difference in contraction fractions at the conclusion of *Stentor* training may be a possible confounder. In the future, it will be useful to explore different training lengths to determine if rate sensitivity emerges in *Stentor* at longer or shorter timescales.

### Hidden variables and temporal response

A major difference between the present model and our previous two-state model is that in the previous model, the states were directly observable based on the frequency of response. In the present model, there are two hidden variables, meaning that they cannot be directly observed in our habituation experiments: the total number of receptors, and the number of receptors that have been internalized. These could obviously be measured if we knew the receptor identity and had molecular tools to detect how much of the receptor was present at the surface and internally, but they are not directly observable in our current stimulus-response data, so from that perspective we consider them hidden variables. Both of these can potentially influence the dynamics of observed behaviors. The fact that receptor internalization can occur during high force stimulation explains the experimental results of **Figure 4C** and **4E**, and the fact that receptor number can drop by degradation explains the subliminal accumulation results of **Figure 5** as well as the potentiation effects of **Figure 6**. It is important to note that the current model is extremely simplified, and that if more detail were to be added, such as different modification states of receptors or different endosome maturation states, the additional hidden variables could support even richer sets of behaviors. The goal of the present study was to determine how many features of *Stentor* habituation could be accounted for with the simplest possible model, as a guide for future experiments and further model development.

### Molecular implementations of the receptor inactivation model

Our model is based on the assumption that the mechanoreceptor that receives the mechanical stimulus, or some element downstream of the receptor, has some probability of becoming reversibly inactivated, and then in the inactive state, can either be re-activated or destroyed. The reversible inactivation could in principle occur by a variety of mechanisms including endocytic internalization, conformational change, or post-translational modifications such as phosphorylation. In the model presented here, we represented internalization as the main cause of response decrement, but any short-term, reversible transition to an inactive state, which can then lead to a further degradation or return back to the active state, would yield essentially the same model. In this sense, the two-state model can be seen to be making an appearance again, but this time on a per-receptor basis, thus not requiring the synchronization of state switching across an entire cell, which was one limitation of our previous two-state phenomenological model (Rajan 2023a).

Determining which of these types of inactivation, whether it be internalization, receptor conformational change to a transient inactive state, reversible phosphorylation, etc., may be at play during habituation in *Stentor*, is an interesting challenge for future work, and we propose two broad types of strategies for pursuing this question. One approach would be an unbiased method in which various "omics" methods, like proteomics and phosphoproteomics, could be brought to bear to identify molecular changes that take place as the cell becomes habituated. Based on our model, relevant modifications would be those for which the time scale of the modification matches the time scale of habituation, and for which the time scale at which the modification reverts back to baseline matches the time scale of forgetting. Such an approach could be successful if the inactivating modification is something that can be detected by appropriate omics methods. But some forms of inactivation, such as internalization into the cell interior to remove the receptor from the plasma membrane, would not be revealed by standard proteomic analyses of whole cells. The alternative approach would to be guess the identity of the receptor, and then directly test whether it is subject to any identifiable modifications or relocalization within the cell that might lead to its inactivation or degradation. At present, this approach seems less promising since it is not obvious what protein plays the role of the mechanoreceptor. In any case we believe that this is an area that is ripe for experimental investigation using the tools of molecular cell biology.

We also note that other *Stentor* behaviors might employ a similar mechanism for decision making. The related *Stentor* species *Stentor roeseli* has multiple possible evasion mechanisms beyond just contraction, including bending and ciliary reversal, and appears to try them out in a pre-defined sequence (Jennings 1902; Jennings 1906; Dexter 2019; Trinh 2019; Marshall 2019). If the different possible responses are triggered by a single underlying receptor molecule as it passes distinct thresholds, this could potentially explain their sequential deployment in a predictable order. Such a model would make two predictions. First, the time spent in the earlier behaviors (bending, reversal) should decrease over repeated rounds of training such that the cell will move more rapidly to the later behaviors (contraction, detachment). This effect was reported by Jennings (1902; 1906), and while not explicitly reported in Dexter (2019), this trend is clearly seen in their data (Marshall 2019). Second, if a single underlying molecule such as the receptor for the noxious stimulus is being inactivated/degraded over time during the procedure, then when the stimulus is removed, the receptor would be re-synthesized such that if a stimulus were to be re-applied at various time-points during the re-synthesis period, the evasion behaviors would be expected to be deployed in the opposite order from when the cell was initially trained. To our knowledge, this experiment has not been done.

### What do cells learn? A kernel estimation framework for Stentor habituation

We have previously tended to consider the habituation process as one in which the cell learns something about the stimulus - deciding if it is caused by a threatening or non-threatening condition in its environment. For *Stentor*, the cell needs to make a quick decision, when it comes into contact with a moving object, whether or not to trigger its contraction response. Contraction is rapid and effective as an except mechanism, but energetically costly. The contraction process itself does not use ATP, just calcium binding to centrin family proteins (Huang and Pitelka 1973; Maloney 2005) but afterwards, it is necessary to pump all that calcium back out, and that incurs a huge energetic cost. Certainly, if a cell is struck with a very strong force that might be typical of a larger predator, contracting would be a good decision. For a sufficiently strong force, it might always make sense to contract since even though this burns energy, at least the cell is not consumed by a predator. A weaker force might be less obviously threatening, but it could still be generated by a large and threatening predator, if the initial contact was not head-on. The only way to know if a weak stimulus is "typical" would be to test whether multiple such stimuli had been received. In the absence of any stimuli, the cell would have no information about the dangers that might be surrounding it, like a child hiding in a dark forest. Therefore, an effective strategy would be to stop contracting if weak stimuli were being detected at some baseline rate, but if a long period goes by without any stimuli, even weak ones, it could be an effective strategy to initially contract when the next stimulus arrives, even if weak, until enough stimuli have been received to make a determination about how typical the weak stimulus is.

In this view, the cell is carrying out an estimation process using impulse data that arrives at random times, and from these randomly arriving inputs, it needs to update an internal representation of the environment that is continuously available for decision making. A classical way to handle such an estimation problem is with kernel estimators, where each impulse is convolved with an impulse response that extends over time, allowing inputs to be integrated over some window, with gradually reducing influence from inputs that recede into the past. In our model, each stimulus drives the internalization of some set of receptors which remain internalized for a random time. In this case, if we ignore degradation and allow only recycling, the kernel is a negative square pulse with exponentially distributed duration. Integration is carried out by the combined effect of all receptor-linked channels on membrane potential.

A recent theoretical study has shown how habituation in general may be cast in the framework of linear filters (Gershman 2024). Our simulations of habituation extent versus stimulus frequency (**Figure 3E**) shows that the cell is acting like a low-pass filter, transmitting a pulse train of input stimuli into a pulse train of output stimuli with a gain that decreases as frequency increases. It would be potentially interesting to explore how the parameters of the current bottom-up molecular model may relate to the statistical framework of Gershman’s linear filter model.

There has been a long discussion in psychology and in philosophy of mind about the definition of learning. Our model represents an example in which the term "learning" is best construed as indicating a process in which an organism updates the parameters of an internal model of the world. Here, the number of receptors serves as a proxy for how safe the environment is, with high numbers of receptors indicating that either there is a definitive threat nearby, or else there is not enough information to know if there is a threat. Often when discussing habituation in a single-cell context, there is a debate about whether habituation is really "just" adaptation. In a multicellular organism, the distinction is based on where the habituation occurs - if it occurs at the sensory or motor levels, it is called adaptation, whereas if it occurs centrally, i.e. within the nervous system, it is called habituation (Staddon and Higa 1996; Kuenen 1981, Torre 1995). The distinction thus has nothing to do with what the process is, or how it is implemented, but rather with how it is used by the organism. In the case of *Stentor*, given that the receptor number is reflecting information about the world, on the basis of which the cell can make a decision about future responses, it does seem that habituation is indeed the more apt description.

## Methods

### Mathematical model for habituation based on mechanoreceptor degradation

In our model, we assume that the stimulus is sensed by some number of mechanoreceptors in the cell membrane that are linked to ion channels, which open with a probability that is a function of the stimulus magnitude. If enough receptor linked channels open so that the change in voltage crosses a threshold, then an action potential fires. Meanwhile, every receptor that is open has a defined probability of being internalized. Once internalized, receptors have a defined probability per unit time of being recycled back to the surface or destroyed via proteolytic degradation. We assume that receptors are continuously synthesized at a constant rate, and receptors are degraded at a constant rate due to normal protein turnover.

We note that while this model is based on known biochemical functions that are widespread in cell biology (receptor internalization, membrane potential, thresholded action potentials), it is still a highly simplistic model that leaves out many biochemical and molecular details. Model parameters are not based on direct experimental information from *Stentor* cells, for which such information is currently lacking. As such, the results of this model are not intended to be directly compared with numerical results from experiments. Instead, our goal is only to ask whether the model demonstrates key qualitative features of *Stentor* habituation that can be observed in experiments. With that in mind, we present the individual components of the model, each focused on a different biochemical process:

### Modeling channel protein dynamics

The number of surface and internalized receptors, as a function of time, are denoted S(t) and I(t). We represent the continuous change in these two receptor pools by the following equations, where k_deg_ describes normal protein degradation and k_des_ describes active destruction of internalized receptors.

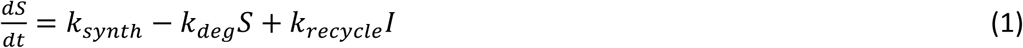

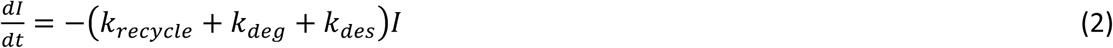

### Modeling mechanoreceptor opening

We represent the stimulus input as a series of delta functions of amplitude F, and assess channel opening only at those timepoints at which one of the stimuli in the series are being delivered. Given a stimulus F, we describe the probability of a mechanoreceptor channel opening by the following equation:

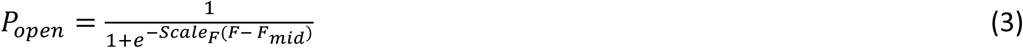

Where F_mid_ represents the stimulus magnitude at 50% maximum activation.

Given P_open_ and the total number of receptors on the surface S(t), we calculate the actual number of receptors that open in response to a given stimulus, Nopen, using Poisson statistics with parameter P_open_S(t).

### Membrane electrical model

Once the number of open channels is determined, we next calculate the effect on membrane potential. We model the membrane as a current source (reflecting ion pumps) in series with a resistor which incorporates the basal membrane resistance combined with any open membrane channels. With no receptor channels open, we assume some membrane conductance Sa. Given the membrane current source J, the basal membrane voltage, which we denote V_1_, is given by

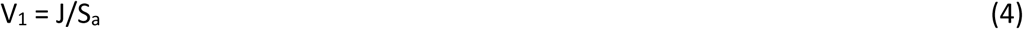

During a stimulus, some number n=Nopen channels open, each with its own conductance S_b_. The net conductance of the membrane thus changes to S_a_ + nS_b_. Now, the overall membrane voltage will become

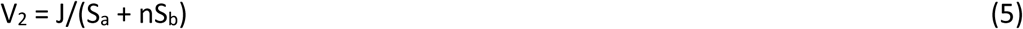

We model the action potential as occurring when the change in membrane voltage ΔV exceeds some threshold V_thresh_. We calculate ΔV as follows

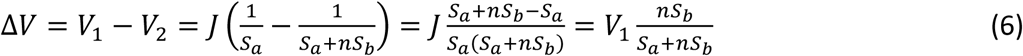

Because both V_1_ and S_b_ are unknown parameters, we introduce the parameter V_i_ = V_1_S_b_ which yields the form of the equation used within the simulation program. When S_a_ >> nS_b_, the change in membrane voltage is approximately proportional to the number of open channels.

### Modeling channel internalization

After channels open, internalization of open channels on the surface then takes place with a probability k_int_, with k_int_*N_open_ determining the number of internalized channels, which is added to I(t) and subtracted from S(t) at the moment of the stimulus. This is a discrete time change that only happens when each stimulus arrives. In between stimulus arrivals, all receptor dynamics are governed by equations 1 and 2. Once internalized, receptors can be recycled back to the surface or removed from the population according to equation 2.

### Stochastic simulation

Using the equations above, we simulate the system using the Euler method with a time step of 0.01. We find a small (less than 10%) discrepancy in the response probabilities between the stochastic and deterministic versions of the model, described below, which is attributable to quantization effects on the receptor number when the average number is small.

### Deterministic approximation

The stochastic model presented above is slow to run, making it difficult to explore various combinations of parameters and input sequences. We therefore developed a deterministic approximation in which we represent S and I by real numbers.

First, we calculate the initial number of receptors by assuming a steady state condition in the absence of any stimuli, such that all receptors are on the surface, and the number of receptors is just k_synth_/k_deg_ as obtained by solving equation 1 for steady state. Next, we consider the stimuli in sequence and, starting with the first stimulus, calculate the probability of any given receptor becoming activated (open) using equation 3. Among the newly activated receptors a fraction is internalized based on the parameter k_int_. The values of surface and internalized receptors are then updated. Next, the minimum number of receptors that would need to be open in order to cross the activation threshold is calculated by solving equation 6 for n. Using the Poisson CDF, we then calculate the probability that enough receptors are activated to equal or exceed this minimal number, given the total number of surface receptors and the probability of each receptor being open. This value gives us the response probability which is the output of the model. Finally, we integrate equations 1 and 2 to determine the new values of S and I at the timepoint when the next stimulus will arrive. The whole procedure is then repeated using these updated values, and this process is repeated for every timepoint. To simulate the forgetting phase after cessation of stimuli, a series of virtual stimuli are modeled such that at each virtual stimulus timepoint, the response probability is calculated as above, but the activation status of channels is not changed, such that S and I are allowed to evolve according to equations 1 and 2 with no effect from the virtual stimuli.

The receptor inactivation model has a large number of parameters, for which experimental values are not yet available. In order to ask whether the model can in principle explain observed behavior, we need a way to choose parameter combinations that produce qualitatively reasonable outputs from the model. In order to obtain parameter values for the simulation, we started with the results of an earlier model, described in Supplemental Materials, which only modeled receptor inactivation but not internalization or recycling. This allowed us to find values for the input/output and dynamics parameters for most of the system, in order to obtain habituation rates and response probabilities that approximated those seen in actual cells. Starting with those parameters, we then adjusted the additional parameters for recycling (k_int_ and kr_ecycle_) as well as re-adjusting the input/output midpoint parameter Fmid (see equation 3).

The result of this parameter search process yielded the following model parameters that were used for all simulations shown in this paper:

k_int_ 0.1
k_recycle_ 0.1
k_synth_ 0.7
k_deg_ 0.02
k_des_ 0.005
F_mid_ 1.5
Scale_F_ 0.6
S_a_ 1000
S_b_ 0.00025
action_threshold 0.012

We note that this parameter search was not an attempt to determine actual parameter values by fitting, but merely an effort to show that the model can, in principle, produce similar behavior to actual cells. We make no claims with respect to uniqueness or optimality of this parameter set - our results do not by any means preclude other parameter combinations giving equal or better performance.

### Stentor cell culture

*Stentor coeruleus* cells, originally obtained from VWR (Ward’s Science 470176-586; Radnor, PA), were grown using a modified version of a previously described culturing protocol (Lin 2018). Cells were grown in a 946-mL glass bowl (Pyrex 1075428; Downers Grove, IL) filled halfway with pasteurized spring water (Carolina Biological 132458; Burlington, NC). Each culture was started with approximately 200-500 cells transferred into a new bowl with fresh media.

When not in use, *Stentor* cultures were stored in a dark drawer at room temperature. Cells used for habituation experiments came from cultures that were 2.5-6 weeks old. To the extent possible, cells for habituation experiments were chosen such that they had a healthy trumpet shape when fully extended (**Figure 1B**) and did not show any preliminary signs of entering mitosis. For any given set of experimental comparisons, cells from both the experimental and control groups were sourced from the same culture. The rate sensitivity experiments, however, were an exception in that the final replicate of the group of cells trained with a frequency of 1 tap per 1.2 seconds was sourced from a different culture due to contamination of the original culture.

Solid cultures of *Chlamydomonas reinhardtii*, grown on Tris-Acetate-Phosphate (TAP) agar plates under bright light, were used as the food source for *Stentor* cultures. Sterile inoculating loops (BD Difco 220217; Franklin Lakes, NJ) were used to scrape approximately 9-15 mg of solid *Chlamydomonas* culture from a TAP plate. The *Chlamydomonas* bolus was dislodged into the *Stentor* culture through manual stirring and shaking of the sterile loop. The bolus was dispersed in the culture by gently pipetting up and down with a P1000 pipette. Remaining large, insoluble fragments of the *Chlamydomonas* bolus were subsequently removed from the *Stentor* culture using the same P1000 pipette. Media removed during this cleaning process was replenished with fresh pasteurized spring water. When experiments were underway, *Stentor* were fed approximately every 1-2 days, depending on culture density. Prior to any experiments on a given day, *Stentor* were examined under dark field illumination to confirm the presence of *Chlamydomonas* inside *Stentor* food vacuoles, thus ensuring that the *Stentor* cells were consistently well fed prior to experimentation.

### Experimental analysis of Stentor habituation to test model predictions

*Stentor* habituation experiments were carried out at room temperature using the protocol described in Rajan et al. 2023b. Following an acclimation period of approximately two hours, mechanical taps were delivered to the cells at a specified force and frequency using an Arduino-controlled habituation device (Rajan 2023b). This device applies mechanical stimulation by striking a metal strip, to which the *Stentor* culture dish is attached, with an aluminum armature driven by a stepper motor. *Stentor* behavior over the course of the habituation experiments was recorded on a USB microscope (Celestron 44308; Torrance, CA) which was positioned over the petri plate such that the maximum number of anchored cells was included in the field of view. A piece of standard white printer paper (22 cm x 28 cm) was placed on top of the USB microscope to soften the ambient glare and to keep the experimental light level at approximately 41 lux. The fraction of cells contracted at various time points during training was subsequently quantified.

For experiments testing the ability of *Stentor* to habituate to low-force stimuli while experiencing high-force stimuli, the cells were given 19 level 3 (high) or level 2 (higher-force) stimuli and subsequently given 41 level 4 (low-force) stimuli, all at a frequency of 1 stimulus per minute. The device is calibrated to four force levels with 1 being the strongest and 4 the weakest force, with the difference among forces created by the number of steps/microsteps taken by the motor. Details of how the force levels are defined is given in Rajan et al. 2023b. To assess the effects of interleaving high-force and low-force stimuli, the cells were given one tap per minute alternating between level 4 taps and level 3 taps. Cells in the control groups received one level 4 tap per minute.

To test subliminal accumulation, *Stentor* were trained with level 4 stimuli at a frequency of 1 tap per 1.2 seconds and then tested for memory retention after training. One group of cells was trained for 20 minutes and the other was trained for 12 hours overnight. Forgetting was tested by delivering level 4 taps at 1.5, 5, 10, 20, 30, 60, and 90 minutes post-training and then quantifying the fraction of cells contracted at each time point.

The effects of high force training on memory retention were tested by training cells for 12 hours overnight at a frequency of 1 tap per minute with either level 4 or level 3 stimuli. Forgetting was tested by delivering level 4 taps at 2, 5, 10, 20, 30, 60, and 90 minutes post-training. Unlike cells in the other experiments included here, these cells did not experience a two-hour acclimation period prior to training. *Stentor* were trained and tested in pasteurized spring water containing 0.25% dimethyl sulfoxide (DMSO).

Potentiation in *Stentor* was induced by delivering successive training bouts. After an acclimation period of approximately two hours, the cells were trained for 60 minutes with level 4 stimuli at a frequency of 1 tap per minute (bout 1). Following a 30-minute break during which the cells did not receive any mechanical stimulation, an additional 60 stimuli were delivered at a frequency of 1 tap per minute (bout 2). This process continued for a total of four training bouts.

To test rate sensitivity, cells were trained with level 4 stimuli for 12 hours overnight and then tested for memory retention following cessation of training. One group of cells was trained at a frequency of 1 tap per minute while the other was trained at a frequency of 1 tap per 1.2 seconds. Once again, forgetting was tested by delivering level 4 taps at 1.5, 5, 10, 20, 30, 60, and 90 minutes post-training and then quantifying the fraction of cells contracted at each time point.

## Supporting information

Supplemental material - degradation-only model for receptor inactivation

## Acknowledgments

We acknowledge stimulating discussions with students in the UCSF Cellular Cognition course held 2012 - 2016 during our initial efforts to reproduce *Stentor* habituation and study its properties in the lab, as well as with students and faculty in the Physiology Course at the Marine Biological Laboratory in Woods Hole where we continued to develop experiments on *Stentor* learning. We also thank current and former members of the Marshall lab, particularly Mark Slabodnick and Tatyana Makushok, as well as Adam Frost and Steve Beckwith for discussions about this project. This work was supported by NSF grant MCB-2012647, the I2CELL award from the Fondation Fourmentin-Guilbert, and NIH grant R35 GM130327.

