## Supplemental material - degradation-only model for receptor inactivation for "A receptor-inactivation model for single-celled habituation in *Stentor coeruleus*"

***Mathematical model for habituation based on mechanoreceptor degradation***

Prior to developing the current receptor internalization model, we had developed a model based on receptor tagging and degradation, in which we assumed that every receptor that is open has a defined probability of being tagged for degradation (for example by ubiquitination). In this model we assume that receptors are continuously synthesized at a constant rate, that receptors are degraded at a constant rate due to normal protein turnover, and that receptors that have become marked for degradation can also be degraded by an alternative degradation pathway that has a different rate than normal turnover. In this version of the model, we did not represent internalization or recycling. The primary difference in the mathematical form of this model from the internalization model is that once a receptor is marked, there is no process for reversing the mark, so eventually all marked receptors will be degraded. Here we described the implementation of this model and the approach for identifying parameters.

***Modeling channel protein dynamics***

The number of unmarked and marked receptors, as a function of time, are denoted U(t) and M(t). We represent the change in these two receptor pools by the following equations, where kdeg describes normal protein degradation and kdes describes active destruction of marked receptors.

$$\frac{dU}{dt} = k_{synth} - k_{deg}U \qquad (1)$$

$$\frac{dM}{dt} = -\left(k_{deg} + k_{des}\right)M \qquad (2)$$

Receptor activation follows the same equation (main text equation 3) as for our current internalization model.

Given Popen and the total number of receptors U(t)+M(t), we calculate the actual number of receptors that open in response to a given stimulus, Nopen, using Poisson statistics with parameter Popen*[U(t)+M(t)]

***Membrane electrical model***

Once the number of open channels is determined, we next calculate the effect on membrane potential. We model the membrane as a current source (reflecting ion pumps) in series with a resistor which incorporates the basal membrane resistance combined with any open membrane channels. With no receptor channels open, we assume some membrane conductance Sa. Given the membrane current source I, the basal membrane voltage, which we denote Vi, is given by

V1 = I/Sa (4)

During a stimulus, some number n=Nopen channels open, each with its own conductance Sb. The net conductance of the membrane thus changes to Sa + nSb. Now, the overall membrane voltage will become

V2 = I/(Sa + nSb) (5)

We model the action potential as occurring when the change in membrane voltage ΔV exceeds some threshold $V_{thresh}$. We calculate ΔV as follows

$$\Delta V = V_1 - V_2 = I\left(\frac{1}{S_a} - \frac{1}{S_a+nS_b}\right) = I\,\frac{S_a+nS_b-S_a}{S_a(S_a+nS_b)} = V_1\,\frac{nS_b}{S_a+nS_b} \tag{6}$$

Because both V1 and Sb are unknown parameters, we introduce the parameter Vi = V1Sb which yields the form of the equation used within the simulation program. When Sa >> nSb, the change in membrane voltage is approximately proportional to the number of open channels.

***Modeling channel marking for degradation***
After channels open, marking of channels for inactivation then takes place with a probability Ptag, with Ptag*U(t) determining the number of newly marked channels, which is added to M(t) and subtracted from U(t). These marked channels are then removed from the population according to the dynamics described above under Modeling Receptor Protein Dynamics.

***Deterministic approximation***
The stochastic model presented above is slow to run, making it difficult to explore various combinations of parameters and input sequences. We therefore developed a deterministic approximation in which we represent U and M by real numbers.

First, we calculate initial the number of receptors by assuming a steady state condition in the absence of any stimuli, such that all receptors are unmarked, and the number of receptors is just ksynth/kdeg as obtained by solving equation 1 for steady state. Next, we calculate the probability of any given receptor becoming activated (open) using equation 3. Among the newly activated receptors a fraction is marked based on the parameter p_tag. The values of marked and unmarked receptors are then updated. Next, the minimum number of receptors that would need to be open in order to cross the activation threshold is calculated by solving equation 6 for n. Using the Poisson CDF, we then calculate the probability that enough receptors are activated, given the total number and probability of each receptor being open, to equal or exceed this minimal number. This value gives us the response probability which is the output of the model. Finally, we integrate equations 1 and 2 to determine the new values of U and M at the timepoint when the next stimulus will arrive. The whole procedure is then repeated using these updated values, and this process is repeated for every timepoint. To simulate the forgetting phase after cessation of stimuli, a series of virtual stimuli are modeled such that at each virtual stimulus timepoint, the response probability is calculated as above, but the activation status of channels is not changed, such that U and M are allowed to evolve according to equations 1 and 2 with no effect from the virtual stimuli.

***Parameter Sweep***

The receptor inactivation model has a large number of parameters, for which experimental values are not yet available. In order to ask whether the model can in principle explain observed behavior, we need a way to choose parameter combinations that produce qualitatively reasonable outputs from the model. It is not computational feasible to use the stochastic simulation for a brute-force evaluation of all possible parameter combinations. Even the deterministic model described above is slow enough that it would make parameter sweeping prohibitively slow. Instead, we developed the following procedure for numerically estimating system output from any given combination of parameters without the need for any simulations. For each parameter combination tested, we calculate the initial response probability, the steady-state (habituated) response probability, the habituation timescale, and the response recovery timescale. These four values are then compared to the desired values, and the parameter combination that minimizes the sum of the errors in all four outputs is taken as the optimal combination. We recognize that other weightings could be used to emphasize particular aspects of the habituation process.

***Estimating response to initial stimulus for parameter sweeping***

We start by directly determining the number of receptors from equation 1 as in the deterministic model. Next we calculate the initial response probability (response to the first stimulus applied a cell that had not previously been exposed) by calculating the activation probability (p_activation) based on the stimulus magnitude according to equation 3, then multiply the activation probability by the initial number of receptors to obtain the average number of open (activated) receptors.

Next, we use the inverse Poisson CDF to determine the minimum number of active receptors that would be observed with a probability equal to the target initial response probability, given the average number of activated receptors.

From that number, we calculate the activation threshold from equation 6, assuming that the number of receptors calculated from the Poisson inverse CDF are active. This calculation directly gives the action threshold parameter Vthresh, which therefore does not need to be swept.

***Estimating Steady-state response for parameter sweeping***

Once we have determined the initial response probability, we next determine the steady state probability after an infinite number of stimuli. Since the stimulus strength is the same as for the initial stimulus, p-activation is the same. The parameter action_threshold is the same for the whole simulation, so it applies here as well. Based on the conductance equation, the action_threshold parameter determines the number nmin of receptors that need to be active to trigger a response. So, if we know how many receptors there are at any given time point, and p_activation, we can determine the probability that nmin of them will be activated.

*Representation of cell state*

Although all we care about in terms of the probability of contraction is the total number of receptors, in order to calculate changes in receptor number over time we need to recognize that marked and unmarked receptors decay at different rates, hence it is more convenient to keep track of both separately, and then add them up when we need to determine contraction probabilities. We thus represent cell state with two variables the number of marked and unmarked receptors.

*Determining steady-state contraction probability*

The basic approach for determining the steady state contraction probability is to first determine the steady-state number of marked and unmarked receptors, based on the steady state condition for each number, and then use them to get the steady state total receptor number from which the response probability can be easily calculated.

We use the following notation:

M0 = number of marked receptors just after a stimulus has arrived
$M_-$ = number of marked receptors just before a stimulus arrives

U0 = number of unmarked receptors just after a stimulus has arrived
$U_-$ = number of unmarked receptors just before a stimulus arrives

At the moment a stimulus arrives, there are M- and U- marked and unmarked receptors, and then the stimulus determines how many of them will become activated via the parameter p_open according to equation 3. Any marked receptors that open will just stay marked, but any unmarked receptors that are activated will become marked with probability ptag.

Therefore:

$M0 = M_- + ptag*Popen*U_-$ (7)
$U0 = U_- - ptag*Popen*U_-$ (8)

Let $\phi$ represent ptag*pactivation, then

$M0 = M_- + \phi U_-$ (9)
$U0 = U_- - \phi U_- = (1-\phi)U_-$ (10)

Now, starting with M0 and U0, we let each evolve forward in time according to the differential equations

$$\frac{dU}{dt} = k_{synth} - k_{deg}U \tag{11}$$

$$\frac{dM}{dt} = -(k_{deg} + k_{des})M \tag{12}$$

After a given interval τ between successive stimuli, we have

$$M(\tau) = M_0 e^{-(k_{deg}+k_{des})\tau} \tag{13}$$

$$U(\tau) = \frac{k_{synth}}{k_{deg}} + \left(U_0 - \frac{k_{synth}}{k_{deg}}\right) e^{-k_{deg}\tau} \tag{14}$$

At steady state, M- = M(τ)=Mss and U- = U(τ)=Uss

Let θ=ksynth/kdeg

We can then solve for Uss as follows:

$$Uss = \theta + (U_0 - \theta)e^{-k_{deg}\tau} \tag{15}$$

$$Uss = \theta + \left((1 - \emptyset)Uss - \theta\right)e^{-k_{deg}\tau} \tag{16}$$

$$Uss - (1 - \emptyset)Usse^{-k_{deg}\tau} = \theta\left(1 - e^{-k_{deg}\tau}\right) \tag{17}$$

$$Uss\left[1 - (1 - \emptyset)e^{-k_{deg}\tau}\right] = \theta\left(1 - e^{-k_{deg}\tau}\right) \tag{18}$$

Which gives the steady state number of unmarked receptors:

$$Uss = \frac{\theta\left(1 - e^{-k_{deg}\tau}\right)}{\left[1 - (1-\emptyset)e^{-k_{deg}\tau}\right]} \tag{19}$$

Now we can plug this into equations 9 and 13:

$$Mss = (Mss + \emptyset Uss)e^{-(k_{deg}+ k_{des})\tau} \tag{20}$$

$$Mss\left[1 - e^{-(k_{deg}+ k_{des})\tau}\right] = \emptyset Usse^{-(k_{deg}+ k_{des})\tau} \tag{21}$$

Which gives the steady state number of marked receptors:

$$Mss = \frac{\emptyset Usse^{-\left(k_{deg}+\,k_{des}\right)\tau}}{\left[1-\,e^{-\left(k_{deg}+\,k_{des}\right)\tau}\right]} \qquad (22)$$

From equations 19 and 22, we obtain the total number of receptors at steady state, Nss = Mss + Uss, From which we can calculate the number that activate in response to the stimulus based on $P_{open}$ (which was determined by the stimulus strength). Finally, we use the Poisson CDF to determine the probability that, given this average number of activated receptors, a sufficient number (nmin, calculated as above) are activated to cross the action threshold. This yields the steady state contraction probability.

***Estimating Learning timescale for parameter sweeping***

Using the initial contraction probability and the steady state contraction probability that have been calculated, we determine the halfway point between the two probabilities. We now seek to determine what the receptor number would be needed to attain that halfway-point response probability. To do this, once n_min, the required number of receptors that need to open in order to cross the activation threshold, has been calculated, we generate a lookup table that contains the Poisson parameter necessary for a Poisson process such that the number of counts will exceed n_min as a function of the desired probability. The Poisson parameter will then equal n_receptor*Popen, so given Popen(which only depends on the stimulus magnitude and the parameters of equation 3), we can determine n_receptor at the halfway response point. Next we calculate the time required to reach this point. Assuming exponential decay of the receptor number from its initial value, we calculate the remaining number of receptors after the first stimulus interval, i.e. the number that will be present at the start of the next stimulus. From this, we take the ratio ρ of receptor numbers after and before the first interval. Assuming exponential decay, this decay ratio should approximately hold for all intervals, and we use this to estimate the number of stimuli required to pass the halfway point.

Defining $N_{init}$ as the initial receptor number prior to any stimuli, and $N_H$ has the number of receptors at the half response point, the number of stimuli take to reach the half response will be

$$Steps = \frac{\ln\, N_H - \ln N_{init}}{ln\rho} \qquad (23)$$

The learning half-time τlearn is thus given by:

τlearn = Steps*stimulus_interval (24)

***Estimating Forgetting timescale for parameter sweeping***

To calculate the forgetting lifetime, we recognize that at the end of the last stimulus period, the number of marked and unmarked receptors will be Mss and Uss. No more marking occurs during the forgetting period, since no stimuli are applied, thus each of these two quantities will undergo exponential decay with its own rate. The total number of receptors will thus decay with an overall decay rate given by the same formula used to calculate fluorescence lifetime of a mixture of two fluorophores (Lakowicz 1983, eq 3.53).

Letting $\alpha$M = Mss/(Mss+Uss) and $\alpha$U = Uss/(Mss+Uss)

And defining the average receptor lifetimes

$$\tau_M = \frac{1}{k_{deg}+k_{des}} \qquad (25)$$

And

$$\tau_U = \frac{1}{k_{deg}} \qquad (26)$$

And using the formulate for average lifetime we get

$$\langle\tau\rangle = \frac{\alpha_M \tau_M^2 + \alpha_U \tau_U^2}{\alpha_M \tau_M + \alpha_U \tau_U} \qquad (27)$$

This is the decay time for the receptor number, and the reciprocal is the decay rate constant for receptor number, krecovery = 1/<t>.

With this decay rate in hand, we can calculate the time for the receptor number at steady state to increase to a number that will restore the response probability to halfway between steady state and initial response probability (which is the same number of receptors as calculated above for the learning rate)

$$\tau_{forget} = \frac{ln\left( {(N_H - N_{init})}/{(N_{ss} - N_{init})} \right)}{-k_{recovery}} \qquad (28)$$

***Parameter search procedure***

For the simulation results in this paper, we carried out the sweep parameter procedure with target values given in Table 1.

Table 1. Target output values for parameter sweep

| value | Low force | High force | High frequency |
|---|---|---|---|

| | | | |
|---|---|---|---|
| Initial response | 0.8 | 0.9 | 0.8 |
| Habituated response | 0.3 | 0.85 | 0.3 |
| Learning timescale | 2 | 12 | 0.5 |
| Forgetting timescale | 3 | 12 | 3 |

Using the above procedure, we scanned the following parameters over the following ranges:
Ksynth 4.0 - 8.0 in steps of 0.5
kdeg 0.01 - 0.2 in steps of 0.01
Fmid 0 - 0.01 in steps of 0.00005
ScaleF 0.05 - 1.0 in steps of 0.05
Sb 0.00005 - 0.005 in steps of 0.00005
Ptag 0 - 0.3 in steps of 0.05
kdes 0.3 - 0.5 in steps of 0.05

These ranges were determined by adjustment using iterative runs of the simulation under different ranges until the procedure found values that were not at either end of their range for any parameter.

Two model parameters did not need to be swept since they only occur in combination with other parameters that were being swept:
Vi was set to 1 such that action_threshold was interpreted as a fraction of Vi.
Sa was set to 1000 but Sb can be rescaled for any other value of Sa.

The result of this parameter sweep process yielded the following model parameters that were used for all simulations shown in this paper:
ksynth 7
kdeg 0.2
Fmid 0.01
ScaleF 0.45
Sb 0.00025
ptag 0.1
kdes 0.45
action_threshold 0.017
